# MOSPD2 a new endoplasmic reticulum-lipid droplet tether functioning in LD homeostasis

**DOI:** 10.1101/2022.02.11.479928

**Authors:** Mehdi Zouiouich, Thomas Di Mattia, Arthur Martinet, Julie Eichler, Corinne Wendling, Nario Tomishige, Erwan Grandgirard, Nicolas Fuggetta, Catherine Ramain, Giulia Mizzon, Calvin Dumesnil, Maxime Carpentier, Bernardo Reina-San-Martin, Carole Mathelin, Yannick Schwab, Abdou Rachid Thiam, Toshihide Kobayashi, Guillaume Drin, Catherine Tomasetto, Fabien Alpy

**Author notes:** correspondence to Catherine Tomasetto, IGBMC, 1 rue Laurent Fries, 67400 Illkirch, France; and Fabien Alpy, IGBMC, 1 rue Laurent Fries, 67400 Illkirch, France.

## Abstract

Membrane contact sites between organelles are organized by protein bridges. Among the components of these contacts, the VAP family comprises endoplasmic reticulum (ER)-anchored proteins, such as MOSPD2, functioning as major ER-organelle tethers. MOSPD2 distinguishes itself from the other members of the VAP family by the presence of a CRAL-TRIO domain. In this study, we show that MOSPD2 forms ER-LD contacts thanks to its CRAL-TRIO domain. MOSPD2 ensures the attachment of the ER to LDs through a direct protein-membrane interaction involving an amphipathic helix that has an affinity for lipid packing defects present at the surface of LDs. Remarkably, the absence of MOSPD2 markedly disturbs the assembly of lipid droplets. These data show that MOSPD2, in addition to being a general ER receptor for inter-organelle contacts, possesses an additional tethering activity and is specifically implicated in the biology of LDs via its CRAL-TRIO domain.

## Introduction

The endoplasmic reticulum (ER), a major membrane-bound organelle of eukaryotic cells, ensures diverse functions such as lipid synthesis, protein synthesis and folding, calcium storage, etc… The ER has a network architecture spreading throughout the cytosol, and is in physical contact with the other organelles such as mitochondria, endosomes/lysosomes, autophagic structures, peroxisomes, lipid droplets and the plasma membrane (Wu *et al*, 2018). These contacts, which do not lead to organelle fusion, are named membrane contact sites. They are scaffolded by protein bridges connecting the two membranes and involving protein–membrane and protein–protein interactions (Gatta & Levine, 2017).

Many molecular players involved in the formation of contact sites have been identified in recent years (Di Mattia *et al*, 2020b; Prinz *et al*, 2019; Wu *et al*, 2018). A few of them act as general receptors that recruit a variety of tethering partners and hold a central role in the biology of inter-organelle contacts. The ER-resident VAP (VAMP-associated protein) protein family plays a major role in the formation of contacts between the ER and the other organelles as well as the plasma membrane. This family comprises 6 proteins divided into two subfamilies. The first sub-family comprises VAP-A, VAP-B and MOSPD2 (motile sperm domain-containing protein 2) which are anchored in the ER membrane by a transmembrane helix (Murphy & Levine, 2016; Di Mattia *et al*, 2018). VAP-A, VAP-B and MOSPD2 act as ER receptors binding multiple protein ligands, either cytosolic or localized on the surface of other organelles, to connect them with the ER (Di Mattia *et al*, 2018; Murphy & Levine, 2016). VAP-A, VAP-B and MOSPD2 have an MSP (Major Sperm Protein) domain exposed to the cytosol; this domain hooks proteins that possess a small linear motif named FFAT (two phenylalanines in an acidic tract) (Mikitova & Levine, 2012; Loewen *et al*, 2003; Di Mattia *et al*, 2018). Recently, a novel subfamily comprising MOSPD1 and MOSPD3 was identified; these proteins have an MSP domain that recognizes FFNT (two phenylalanines in a neutral tract) motifs (Cabukusta *et al*, 2020).

The function of VAP-A and VAP-B is well studied, they are central proteins for the formation of inter-organelle contacts and function in lipid transport, ion homeostasis, and autophagy (Murphy & Levine, 2016; Wilhelm *et al*, 2017; Zhao *et al*, 2018; Mesmin *et al*, 2013; Kirmiz *et al*, 2018; Johnson *et al*, 2018). In contrast, the function of MOSPD2 is still elusive. MOSPD2 is a genuine member of the VAP family: it shares with VAP-A and VAP-B a large number of tethering partners and promotes the formation of contacts between the ER and many organelles (Di Mattia *et al*, 2018, 2020a). Unlike VAP-A, VAP-B, MOSPD1, and MOSPD3 which only possess an MSP domain in addition to their TM domain, MOSPD2 possesses an additional cytosolic domain named CRAL-TRIO [cellular retinaldehyde-binding protein (CRALBP) and triple functional domain protein (TRIO)] at its amino terminus. The CRAL-TRIO domain (also called Sec14 domain), present in 28 proteins in human, contains a hydrophobic pocket that in the case of Sec14p and alpha-tocopherol transfer protein (TTPA) can bind/transport small lipophilic molecules such as phospholipids or tocopherols (Chiapparino *et al*, 2016). Knowing that MOSPD2 contains a CRAL-TRIO domain, we hypothesized that it may have a broader function than VAP proteins. Here, we show that in addition to serving as a VAP-like tether that establishes ER-organelle contacts through protein-protein interactions, MOSPD2 also tether the ER to lipid droplets (LDs) by protein-membrane interactions. LDs are ubiquitous organelles that serve as universal lipid stores in cells; they consist of a neutral lipid oil core surrounded by a monolayer of phospholipids associated with peripheral proteins (Olzmann & Carvalho, 2019; Thiam & Beller, 2017; Thiam *et al*, 2013). In this report, we found that the absence of MOSPD2 compromises lipid droplet assembly, showing that MOSPD2 is involved in the biology of LDs.

## Results

### MOSPD2 is involved in the biology of LDs

MOSPD2 is an ER-resident protein (Di Mattia *et al*, 2018). When expressed in HeLa cells, GFP-labeled MOSPD2 exhibited a distinctive reticular localization pattern throughout the cytoplasm, and co-localized with the ER marker Calnexin (Fig. 1A, a). Remarkably, in about half of the cells, MOSPD2 was additionally found in ring- and comma-shaped structures that were also positive for the ER marker Calnexin (Fig. 1A, b). This shows that in some cells, MOSPD2 can be enriched in sub-regions of the ER. Importantly, VAP-A and VAP-B were never observed in similar ring-like structures, suggesting that only MOSPD2 can specifically populate these ER subdomains (Fig. 1B and S1A). Then, we searched whether these MOSPD2-enriched areas corresponded to contact sites between the ER and a particular organelle by performing co-localization experiments using markers of early endosomes (EEA1), late endosomes/lysosomes (Lamp1), mitochondria (OPA1), Golgi (GM130) (Fig. S2 A-D), and LDs (BODIPY 493/503) (Fig. S1B). MOSPD2-positive rings did not overlap with endosomes, Golgi and mitochondria. In contrast, ring- and comma-shaped MOSPD2-positive structures were found to be around LDs labeled with BODIPY 493/503. To confirm this observation, LD biogenesis in HeLa cells was stimulated by oleic acid (OA) treatment (Listenberger & Brown, 2007). After this treatment, most HeLa cells had numerous and large LDs massively surrounded by ring and comma-shaped structures positive for MOSPD2 (Fig. 1C). This accumulation of MOSPD2-positive ER around LDs was also found in other tested cell lines, including hepatocytes (Huh7), melanoma (501-mel), and mammary epithelial (MCF7) cells (Fig. S1C-E).

**Figure 1:**
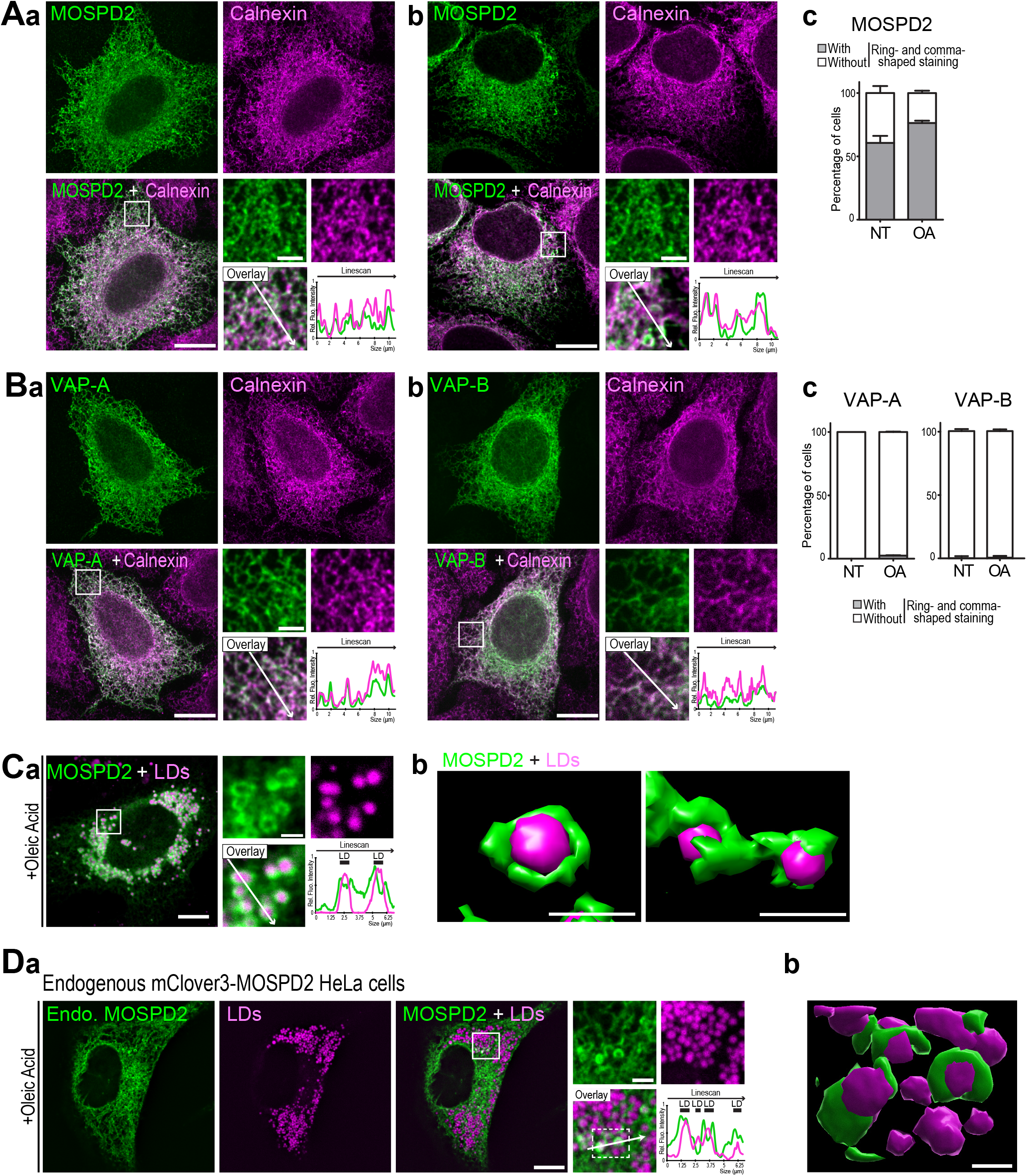
MOSPD2 is an ER-resident protein found enriched around lipid droplets (LDs). A. HeLa cells expressing GFP-MOSPD2 (green) were labeled with an anti-Calnexin antibody (magenta). GFP-MOSPD2 exhibited a reticular pattern (a), with additional ring- and comma-shaped structures (b). Spinning disk confocal microscope (Nikon CSU-X1, 100x NA 1.4) images. Scale bars: 10 μm (insets 2 μm). c: percentage of cells with GFP-positive ring- or comma-shaped structures, in the absence of treatment (NT) or after oleic acid treatment (OA). Mean ± SD; n = 3 independent experiments (NT: 300 cells, OA: 136 cells). B. a: GFP-VAP-A (green) was labeled as in A. Scale bars: 10 μm (insets 2 μm). b: Quantification as in (Ac) of VAP-A and VAP-B expressing cells. Mean ± SD; n = 3 independent experiments (GFP-VAP-A: NT: 119 cells, OA: 126 cells; GFP-VAP-B: NT: 140 cells, OA: 141 cells). C. a: HeLa cells expressing GFP-MOSPD2 were treated with OA (400 μM, overnight) and LDs were labeled with BODIPY 493/503. Confocal microscope (Leica SP5, 63x NA 1.4) images. Scale bars: 10 μm (insets 2 μm). b: 3D reconstruction from confocal images of MOSPD2-positive ER (green) around LDs (magenta). Images generated with the surface representation tool of the Chimera software (Pettersen et al, 2004). Scale bar: 500 nm. D. a: Live imaging of CRISPR/Cas9-edited HeLa cells expressing mClover3-MOSPD2 (green) at the endogenous level, treated with OA, and labeled with LipidTOX (magenta). Images were acquired on a spinning disk confocal microscope (Nikon CSU-X1, 100x NA 1.4). Scale bars: 10 μm (insets 2 μm). b: 3D reconstruction from confocal images of MOSPD2-positive ER (green) around LDs (magenta) using Imaris (white rectangle from overlay panel). Scale bar: 500 nm.

To examine whether endogenous MOSPD2 can be seen in association with LDs, we generated a reporter cell line. The endogenous MOSPD2 gene was modified in HeLa cells using the CRISPR/Cas9 method to fuse the fluorescent protein mClover3 coding sequence in frame with that of MOSPD2 (Fig. S3). As observed by expressing GFP-MOSPD2, endogenous mClover3-MOSPD2 protein was present in structures in contact with LDs (Fig. 1D). Next, in Huh-7 and 501-MEL cells, which express higher levels of endogenous MOSPD2 than HeLa cells (Fig. S3C), we could analyze this protein using a specific antibody (Fig S3 D, E). We found that the endogenous MOSPD2 did accumulate in ring- and comma-shaped structures around LDs in Huh7 and 501-MEL cells treated with OA (Fig. S3 D, E). These data indicated that endogenous MOSPD2 is partially localized around LDs.

These observations prompted us to investigate whether MOSPD2 has a role in LD biology. We first established several cell models of MOSPD2 deficiency. MOSPD2 was either knocked-down using a pool of siRNAs, or knock-out using a CRISPR/Cas9 approach in HeLa cells (Fig. 2A); two independent knock-out clones (KO#1 and KO#2) were analyzed. Then, the number and size of LDs labeled with BODIPY 493/503 were quantified in MOSPD2 knocked-down, knocked-out, and control cells (Fig. 2B, C). Strikingly, compared to control cells, MOSPD2-deficient cells contained less (^~^2-fold) and smaller (^~^40% decrease) LDs. To compare the effect of MOSPD2 deficiency to that of VAP-A and VAP-B, we silenced VAP-A and VAP-B either individually or together using pools of siRNAs (Fig. S5A). Then, LDs were labeled (Fig. S5B) and their number and size quantified (Fig. S5C). The silencing of VAP-A and/or VAP-B had no effect on the number and size of LDs, thus showing that among the FFAT-binding proteins of the VAP family, only MOSPD2 has a specific role in LD biology.

**Figure 2:**
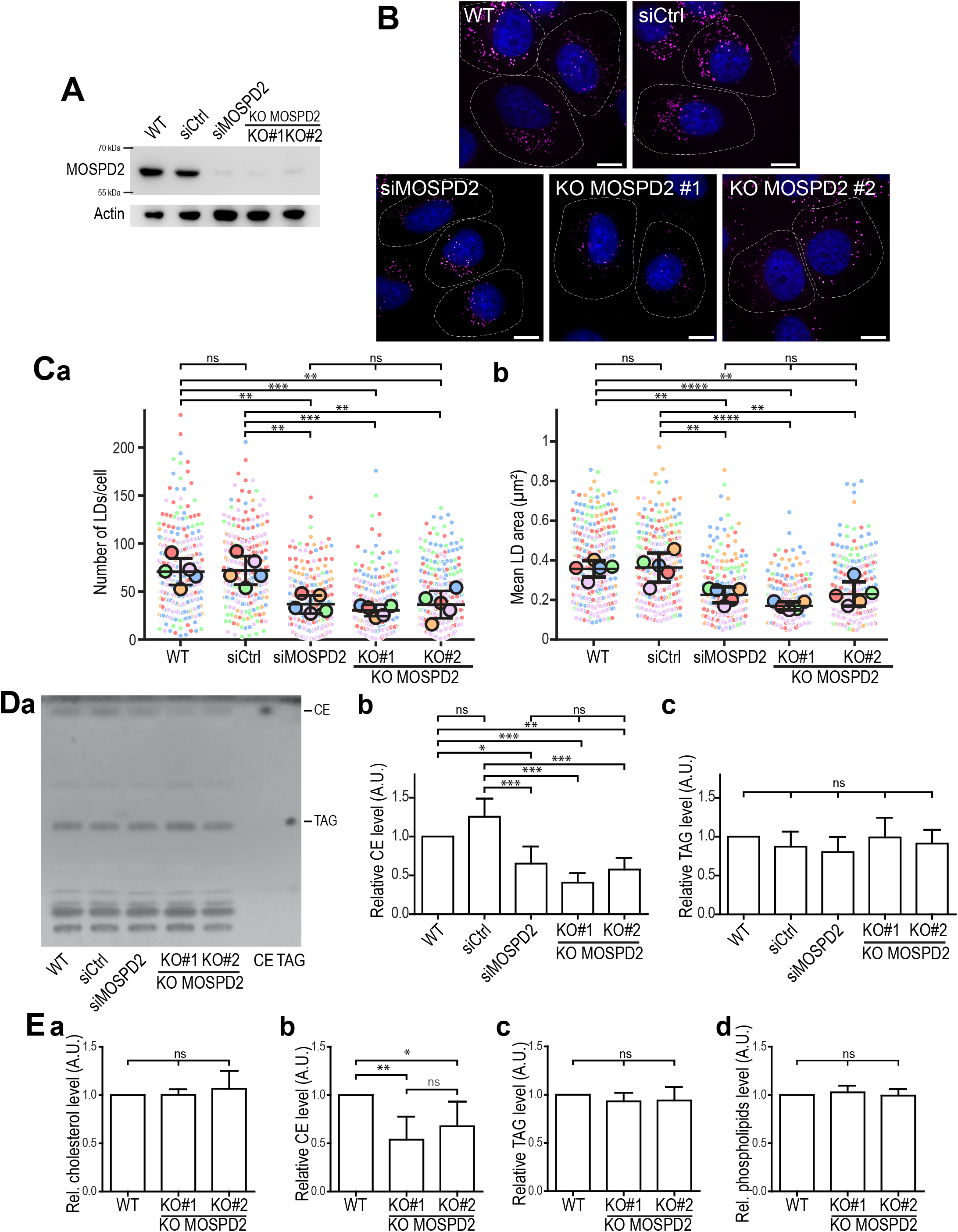
MOSPD2 is involved in LD homeostasis. A. Western blot analysis of MOSPD2 protein level in control HeLa cells (WT), HeLa cells transfected with control siRNAs (siCtrl), and with siRNAs targeting MOSPD2 (siMOSPD2), and in two MOSPD2-deficient HeLa cell clones (KO#1 and KO#2) established by CRISPR/Cas9 gene editing. B. Representative confocal images of parental HeLa cells (WT), of HeLa cells transfected with control siRNAs (siCtrl), and with siRNAs targeting MOSPD2 (siMOSPD2), and of MOSPD2-deficient HeLa cell clones (KO#1 and KO#2) labeled with BODIPY 493/503 (LDs, magenta) and Hoechst 33258 (nuclei, blue). Scale bars: 10 μm. Images were acquired on a spinning disk confocal microscope (Nikon CSU-X1, 100x NA 1.4). C. Number (a) and area (b) of LDs in cells shown in B. Data are displayed as Superplots (Lord et al, 2020) showing the mean number (left) and area (right) of LDs per cell (small dots), and the mean number (left) and area (right) of LDs per independent experiment (large dots). Independent experiments (n=5) are color-coded. Means and error bars (SD) are shown as black bars. Data were collected from 398 (WT), 323 (siCtrl), 280 (siMOSPD2), 333 (KO#1) and 413 (KO#2) cells. One-way ANOVA with Tukey’s multiple comparisons test (**: P < 0.01; ***: P < 0.001; ****: P<0.0001; ns: not significant; n = 5 independent experiments). D. a: Thin layer chromatography (TLC) analysis of lipids extracted from cells shown in A. Neutral lipids were separated with hexane/diethylether/AcOH (80:20:2) and revealed with primuline. Cholesterol ester (CE) and triacylglycerol (TAG) were used as standards. b, c: Relative levels of CE (b) and TAG (c) detected by TLC. Means and error bars (SD) are shown. One-way ANOVA with Tukey’s multiple comparisons test (*: P < 0.05; **: P < 0.01; ***: P < 0.001; n = 4 independent experiments). E. Enzymatic quantification of cholesterol (a), cholesterol ester (b), triglyceride (c) and phospholipid (d) in control HeLa cells (WT) and MOSPD2-deficient HeLa cell clones (KO#1 and KO#2). Means and error bars (SD) are shown. One-way ANOVA with Tukey’s multiple comparisons test (*: P < 0.05; **:P < 0.01; n = 6 independent experiments).

We next determined the level of neutral lipids in MOSPD2-deficient cells, by quantifying total cellular triacylglycerols (TAG) and cholesterol esters (CE) using thin-layer chromatography (TLC) (Fig. 2D). In MOSPD2-deficient cells, TAG levels were unchanged while CE were reduced by ^~^40 %. To further substantiate this phenotype, cholesterol, phospholipids, CE and TAG were quantified by enzymatic methods in wild type and MOSPD2 knock-out cells (Fig. 2E). While cholesterol, phospholipids and TAG remained unchanged in MOSPD2-deficient cells, CE levels were reduced by ^~^40 %. These data show that in the absence of MOSPD2, the level of neutral lipids especially CE is decreased.

Together, these data suggest that MOSPD2 is present in ER sub-domains in contact with LDs and show that MOSPD2 is involved in the biology of LDs.

### MOSPD2 is dynamically recruited to ER-LD contacts

MOSPD2 is present in comma-shaped structures that are also positive for the ER marker Calnexin. This suggests both that MOSPD2 is in ER sheets that do not completely encircle LDs, and that it does not translocate at the surface of LDs, meaning that it would be present in ER-LD contact sites. To examine this, we performed correlative light and electron microscopy (CLEM). GFP-MOSPD2 expressing cells were processed for electron microscopy and embedded in a fluorescence – preserving resin. Thick sections were imaged by spinning disk confocal microscopy and then by transmission electron microscopy (Fig. 3A). GFP-MOSPD2 fluorescence correlated with the presence of ER sheets in contact with LDs. Noteworthy, we confirmed that comma-shaped fluorescent structures corresponded to areas of ER in contact with LDs. Fluorescence was absent from the surface of LDs making no contact with the ER. These data indicate that MOSPD2 accumulates in ER regions in contact with LDs.

**Figure 3:**
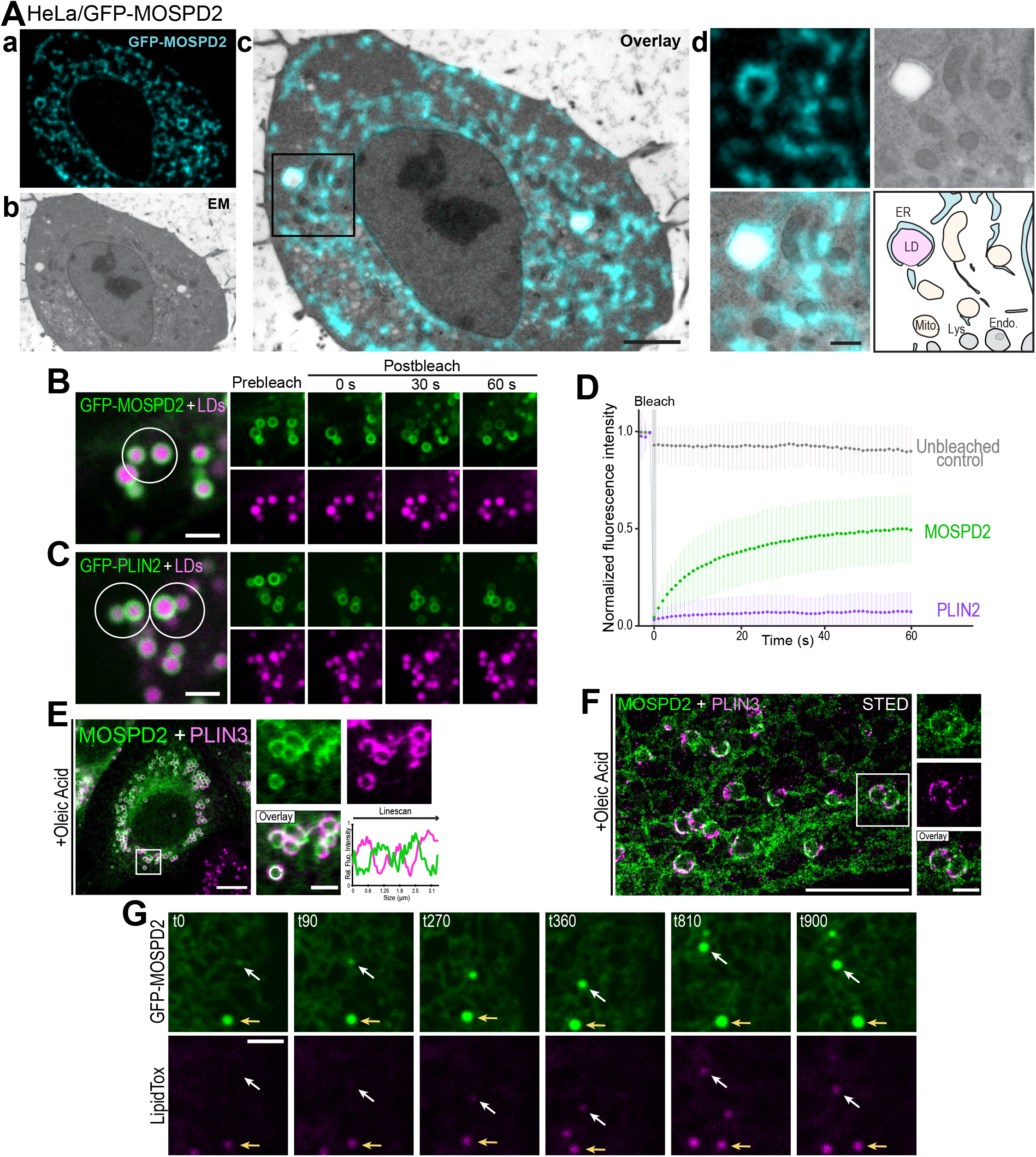
MOSPD2 is dynamically recruited to ER-LD contact sites. A. Correlative light and electron microscopy (EM) of HeLa/GFP-MOSPD2 cells. a: fluorescence microscopy image; b: EM image; c: correlation of GFP-MOSPD2 and EM images (scale bar: 2 μm); d: composite showing higher magnification images of the area outlined in black in (c) (scale bar: 500 nm); bottom right: interpretation scheme showing contacts between organelles; ER and lipid droplets are in cyan and pink, respectively. Mitochondria and endosomes/lysosomes are in yellow and gray, respectively. B-D. FRAP analysis of MOSPD2 and Plin2 mobility. GFP-MOSPD2 (B) and GFP-Plin2 (C) expressing cells were treated with OA and labeled with LipidTOX. GFP fluorescence was photobleached in the area outlined in white, and images acquired every second over a 1-minute period. Scale bars: 2 μm. D: lineplot showing the relative fluorescence intensity in the photobleached region of GFP-MOSPD2 (green) and GFP-Plin2 (blue) expressing cells. The grey curve shows the relative fluorescence intensity of GFP-positive LDs that were not bleached. Means and error bars (SD) of relative fluorescence intensities of 56 (GFP-MOSPD2), 57 (GFP-Plin2) and 72 (unbleached control) regions of interest from 20, 13 and 26 cells, respectively. Data from 3 independent experiments. E, F. HeLa cells expressing GFP-MOSPD2 (green) were treated with OA and labeled with anti-Plin3 antibodies (magenta). Images were acquired by confocal microscopy (Leica SP8) (E), or by STED super-resolution microscopy (F). MOSPD2 and PLIN3 were heterogeneously distributed around LDs. Scale bar: 10 μm (insets 2 μm) in (E) and 5 μm (insets 1 μm) in (F). Subpanels on the right are higher magnification images of the area outlined. The overlay panel shows merged channels. In (E), linescan shows fluorescence intensities of the green and magenta channels along the white circular arrow of the overlay subpanel *i.e*. at the surface of LD). G. HeLa cells expressing GFP-MOSPD2 were imaged live during LDs induction (stained with LipidTOX) by OA addition. The white arrow shows an enrichment of MOSPD2 signal before the appearance of LipidTOX staining. The yellow arrow shows the growth of a LD positive for MOSPD2 before the start of the induction. Images were acquired every 90 seconds (t0-900) on a spinning disk confocal microscope (Nikon CSU-X1, 100x NA 1.4). Scale bar: 2 μm.

That MOSPD2 is both distributed throughout the entire ER membrane and enriched in subdomains of the ER surrounding LDs suggests that it is in equilibrium between ER-LD contact sites and the remainder of the ER. To analyze the dynamics of MOSPD2 movement between these two regions, we performed fluorescence recovery after photobleaching (FRAP) experiments. GFP-MOSPD2 was expressed in HeLa cells and individual LDs were bleached (Fig. 3B and D). GFP-MOSPD2 fluorescence rapidly recovered (t1/2 of ^~^8 s) and reached a plateau equivalent to ^~^50% of the initial fluorescence (Fig. 3B and D). In comparison, the LD protein Perilipin-2 (PLIN2) fused with GFP remained permanently associated with LDs with no recovery observed for the duration of the experiment (Fig. 3C and D). The localization of MOSPD2 is balanced between distinct areas of the ER, some of which being in contact with the LDs, and half of the protein pool can rapidly move in and out these two regions.

We noted that the MOSPD2 signal was not uniform around LDs, suggesting that LDs might have sub-domains. To examine this, we performed co-localization experiments between GFP-MOSPD2 and endogenous PLIN3, a coat protein of LDs. Of interest, while both proteins were present around LDs, they exhibited a different distribution, the two signals being mostly mutually exclusive, as observed by confocal and super-resolution microscopy (Fig. 3E and F). This supports the notion that, in terms of protein composition, the surface of LDs has distinct territories, either sticking to the ER (MOSPD2 territory) or being free (PLIN3 territory) and that contacts define them.

MOSPD2 is recruited around mature LDs. Next, to determine whether MOSPD2 is already recruited early in the life of LDs, we visualized the location of GFP-MOSPD2 following the induction of LD biogenesis by OA by live cell imaging (Fig. 3G). As shown in Fig. 3G, GFP-MOSPD2 was recruited early during LD biogenesis: GFP-MOSPD2 was already present in areas that were not yet detected by the LD marker LipidTox. Thus, MOSPD2 is associated with LDs at different stages of their life.

Overall, these results demonstrate that MOSPD2 dynamically distributes between specific subdomains of the ER: ER membranes in contact with LDs, and the remainder of the ER.

### MOSPD2 tethers the ER to LDs

Because associations between the ER and LDs are frequent events, we wondered whether MOSPD2 merely populates existing ER-LD contacts or, instead, contributes to making these contacts. To address this question, we determined by transmission electron microscopy (TEM) whether the overexpression of MOSPD2 generates more ER-LD contacts. In control cells, LDs made few focal contacts with the ER (Fig. 4A, a). In contrast, in cells expressing MOSPD2, LDs were frequently associated with long segments of ER encircling them (Fig. 4A, b).

**Figure 4:**
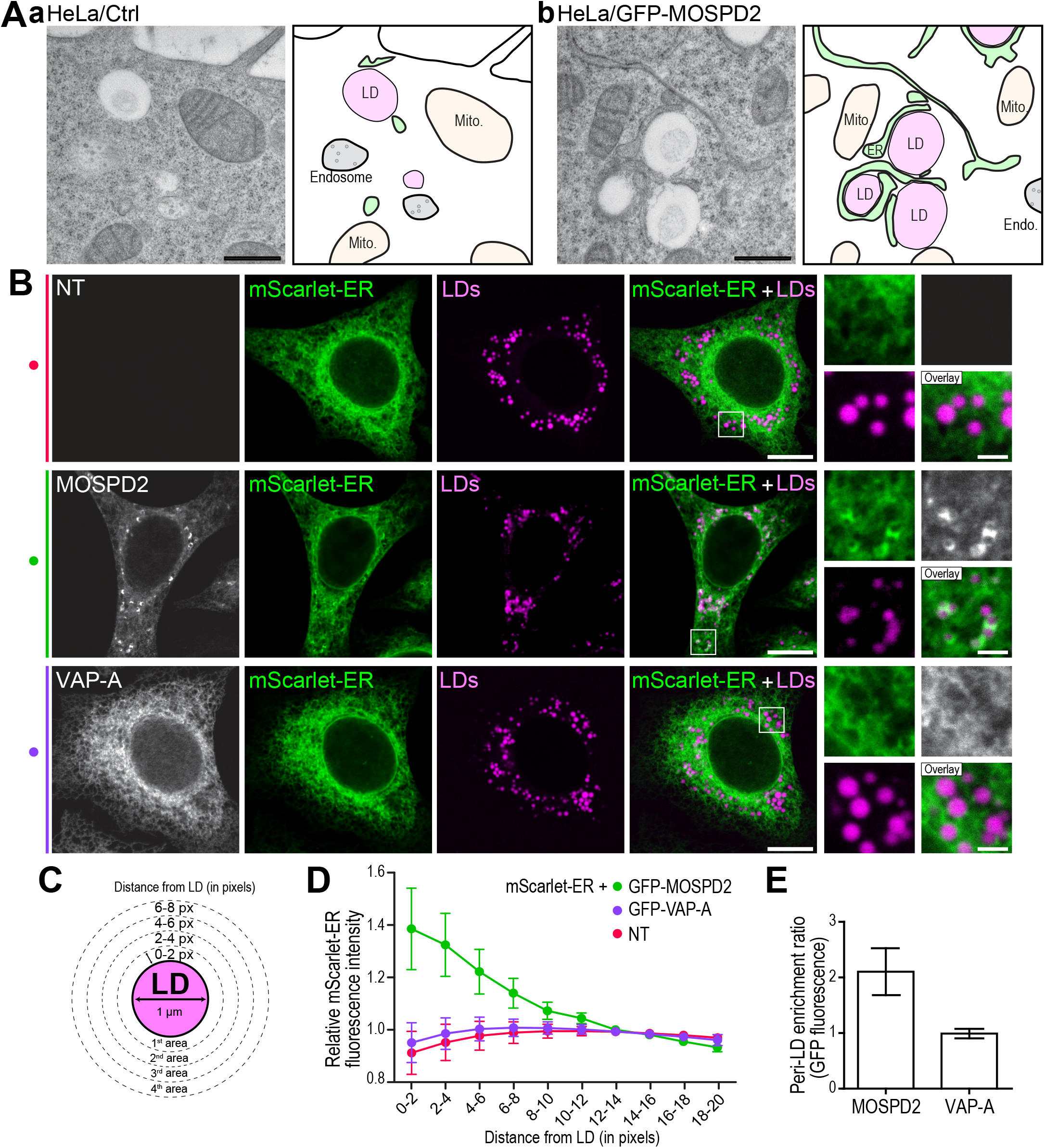
MOSPD2 regulates ER-LD contact sites. A. TEM images of control HeLa (a) and HeLa/GFP-MOSPD2 cells (b) with their interpretation scheme; the ER and LDs are in green and magenta, respectively. Mitochondria and endosomes are in yellow and gray, respectively. Scale bars: 500 nm. B. HeLa cells stably expressing the mScarlet-ER marker (green) were either not transfected (NT, top), transfected with GFP-MOSPD2 (gray, middle), or with GFP-VAP-A (gray, bottom). Cells were treated with oleic acid (50 μM for 6 hours) and LDs stained with LipidTOX (magenta). Images were acquired on a confocal microscope (Leica SP5; x63 NA 1.4). Scale bars: 10 μm (insets 2 μm). C. Schematic representation of the method used for fluorescence quantification around LDs: two pixels-wide areas were segmented around LDs (here represented for a 1 μm wide LD), and the mean mScarlet fluorescence intensity was measured in each area. D. Fluorescence intensity of the ER marker mScarlet-ER around LDs in untransfected (NT, red), GFP-MOSPD2 (green), and GFP-VAP-A (purple) transfected cells. Means ±SD (NT: 39 cells; GFP-MOSPD2: 42 cells; GFP-VAP-A: 46 cells; from 4 independent experiments). The relative mScarlet fluorescence intensity corresponds to the mean fluorescence intensity of mScarlet in each area, divided by the mean fluorescence intensity in the cytoplasm away from LDs (10-20 pixels distance from LDs). E. Relative enrichment of GFP-MOSPD2 and GFP-VAP-A around LDs. The Peri-LD enrichment ratio is the ratio of the mean GFP fluorescence intensity (GFP-MOSPD2 or GFP-VAP-A) in the vicinity of LDs (0-4 pixels distance from LDs; see C), to the mean fluorescence intensity of GFP at a distance from LDs (10-20 pixels distance from LDs). MOSPD2 fluorescence is twice as high around LDs as in the remainder of the cytoplasm, whereas VAP-A fluorescence is at the same level next to LDs and in the rest of the cytoplasm. Means ±SD (GFP-MOSPD2: 42 cells; GFP-VAP-A: 46 cells; data from 4 independent experiments).

To evaluate the capacity of MOSPD2 to drive ER-LD contact formation more quantitatively, we examined by light microscopy the recruitment of the ER around LDs in cells where MOSDP2 was overexpressed. The ER surface was labeled with a red fluorescent marker (mScarlet-ER) and the radial distribution of the fluorescence signal was measured around LDs (Fig. 4B and C, and Fig. S4B). In cells expressing the mScarlet-ER marker alone, the fluorescent signal was evenly distributed in the cytosol, with no enrichment around LDs (Fig. 4B and D). In contrast, in cells expressing MOSPD2, the ER marker accumulated in the periphery of LDs, thus showing that MOSPD2 expression promotes the formation of ER-LD contacts. This was specific to MOSPD2 since VAP-A expression did not result in ER accumulation around the LDs (Fig. 4B and D). Furthermore, consistent with data from Fig. 1, the relative fluorescence of MOSPD2 was 2-fold higher at the periphery of LDs than in the remaining part of the cytoplasm, whereas VAP-A fluorescence was homogeneously distributed; this confirmed that MOSPD2 accumulates around LDs (Fig. 4E).

We next addressed whether MOSPD2 association with LDs relies on Seipin. Seipin, a protein encoded by the *BSCL2* gene which is mutated in lipodystrophy, is a major tether localized in ER-LD junctions tightly controlling LD assembly (Szymanski *et al*, 2007; Salo *et al*, 2016; Gao *et al*, 2019). Since we found MOSPD2 at ER-LD contacts, especially during the early stages of LD formation (Fig.3G), we asked whether this localization relies on Seipin. We found that in cells knock-out or silenced for Seipin, MOSPD2 still localized to ER-LD contacts (Fig. S6), indicating that Seipin was not required for MOSPD2-mediated ER-LD contact formation.

Collectively, these data show that MOSPD2 favors the tethering of the ER with LDs.

### The MSP domain of MOSPD2 is dispensable for the formation of ER-LD contacts

MOSPD2 contains an MSP domain involved in protein-protein interactions, and a CRAL-TRIO domain which is potentially a lipid transfer domain (Di Mattia *et al*, 2018; Chiapparino *et al*, 2016). We first reasoned that the MSP domain could mediate the formation of ER-LD contacts to position the CRAL-TRIO domain at the ER/LD interface. To test this hypothesis, we constructed a deletion mutant lacking the MSP domain (Fig. 5A). This mutant was transfected in HeLa cells in which LD biogenesis was induced by OA, and we analyzed its ability to form ring- and comma-shaped structures around LDs, *i.e*. to form ER-LD contacts. Unexpectedly, we observed that the MOSPD2 ΔMSP mutant localized in ring- and comma-shaped structure around LDs like the WT protein; in fact, we noted that it was even more recruited around LDs than the WT protein (97% vs 77%, Fig. 5B and C). Likewise, the mutation of two key residues in the MSP domain (R404D/L406D referred to here as RD/LD mutant) precluding the recognition of FFAT motifs (Di Mattia *et al*, 2018) resulted in a massive recruitment of MOSPD2 at the periphery of LDs in most cells, both in the presence (Fig. 5A-C) and in the absence of OA (Fig. S7B). Consistent with this observation, CLEM experiments performed with cells expressing GFP-MOSPD2 RD/LD mutant (Fig. S7A) showed that the protein accumulated in ER strands in tight contact with LDs.

**Figure 5:**
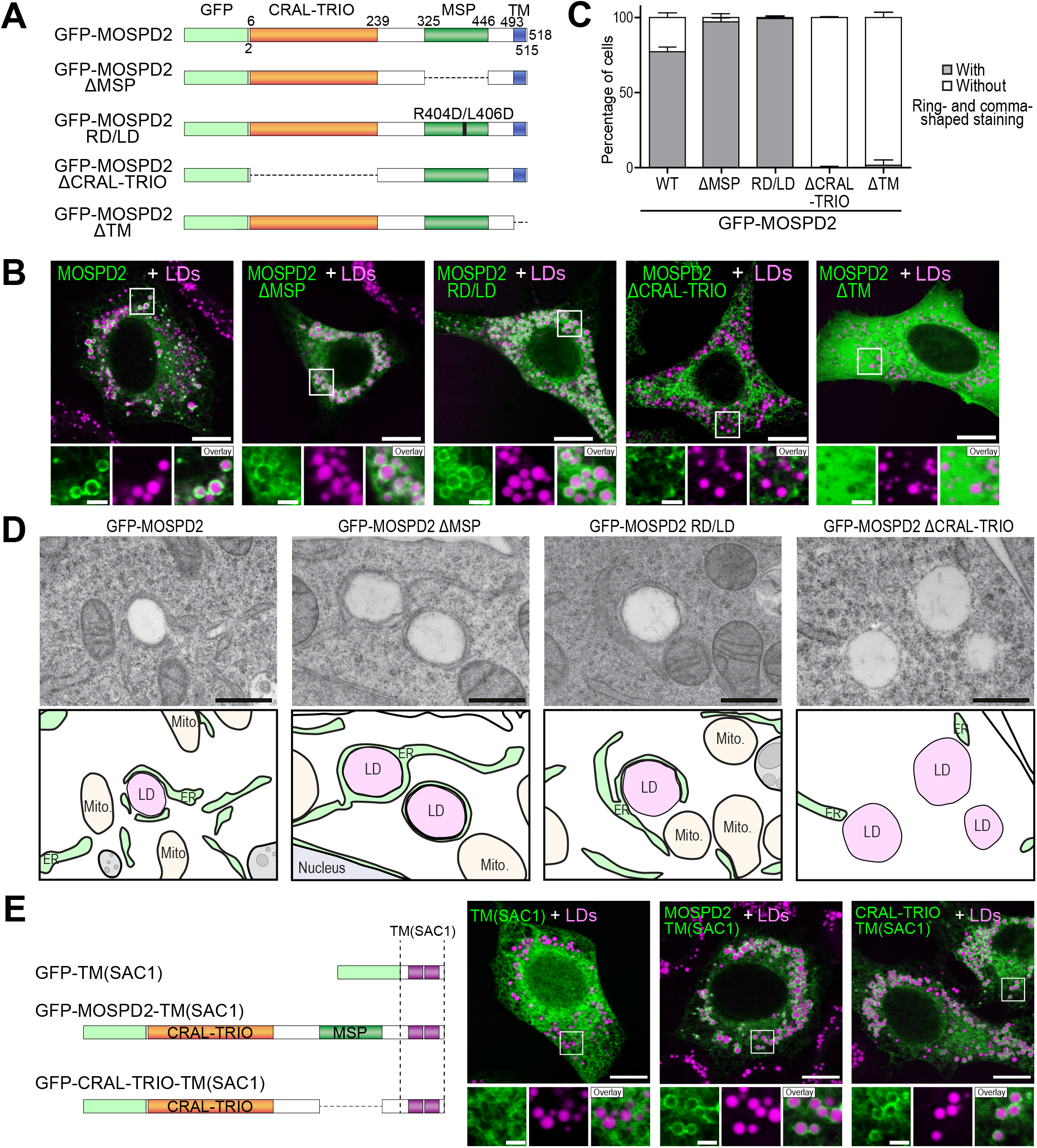
ER-LD contact sites mediated by MOSPD2 depends on its CRAL-TRIO and TM domains. A. Schematic representation of the different WT and mutant proteins used in the study. Two kind of mutants were utilized: deletions of specific domains (ΔCRAL-TRIO, ΔMSP, ΔTM) and point mutation (RD/LD) impairing the MSP domain function. B. Representative confocal images of the GFP-MOSPD2 WT and mutants (green) localization. Cells were treated with OA and LDs stained with Nile Red (magenta). Confocal microscope (Leica SP8; 63x NA 1.4) images. Scale bars: 10 μm (insets 2 μm). C. Quantification of cells presenting ring- and comma-shaped staining. Mean ± SD; n = 3 independent experiments (WT: 67 cells; ΔMSP: 138 cells; RD/LD: 152 cells; ΔCRAL-TRIO: 140 cells; ΔTM: 64 cells). D. EM images of HeLa/GFP-MOSPD2, HeLa/GFP-MOSPD2 ΔMSP, HeLa/GFP-MOSPD2 RD/LD, and HeLa/GFP-MOSPD2 Δ CRAL-TRIO cells (top) and their interpretation scheme (bottom); the ER and LDs are in green and magenta, respectively. Mitochondria and endosomes are in yellow and gray, respectively. Scale bars: 500 nm. E. Left: schematic representation of the different chimeric constructs in which the MOSPD2 TM domain is replaced by the TM of SAC1 (purple). Right: localization of these chimeric proteins (green) and LDs stained with Nile Red (magenta) in HeLa cells treated with OA. Images were acquired on a confocal microscope (Leica SP8; 63x NA 1.4). Scale bars: 10 μm (insets 2 μm). Data information: In (B, E), subpanels on the bottom are higher magnification images of the area outlined. The overlay panel shows merged channels.

To further ascertain that the MSP domain of MOSPD2 is dispensable for the formation of ER-LD contacts, cells expressing GFP-tagged MOSPD2 deletion mutant (GFP-MOSPD2 ΔMSP) and the RD/LD mutant (GFP-MOSPD2 RD/LD) were processed for TEM (Fig. 5D). In cells expressing MOSPD2 mutants lacking the MSP domain (GFP-MOSPD2 ΔMSP), or having a defective MSP domain (GFP-MOSPD2 RD/LD), the ER remained extensively attached to LDs (Fig. 5D).

Thus, the MSP domain is not involved in the formation of ER-LD contacts, and may even limit the recruitment of MOSPD2 at ER-LD contacts.

### The CRAL-TRIO and TM domains of MOSPD2 mediate the formation of ER-LD contacts

Since the MSP domain of MOSPD2 is not necessary for the formation of ER-LD contacts, we alternatively examined the contribution of the TM and CRAL-TRIO domains by testing deletion mutants (GFP-MOSPD2 ΔCRAL-TRIO and GFP-MOSPD2 ΔTM) (Fig. 5A). In contrast to WT MOSPD2 that accumulated in ring- and comma-shaped structures, MOSPD2 devoid of the CRAL-TRIO domain had a reticular-only localization, thus showing that the CRAL-TRIO domain is necessary for the recruitment of MOSPD2 to LDs (Fig. 5B and C).

Cells expressing the GFP-MOSPD2 ΔCRAL-TRIO deletion mutant were further analyzed by TEM. Compared to cells expressing WT MOSPD2 and in which extended and frequent ER-LD contacts were observed, cells expressing this mutant only harbored focal ER-LD contacts (Fig. 5D). Jointly, these results point to a crucial role of the CRAL-TRIO domain for the ability of MOSPD2 to create ER-LD contacts.

Besides, we observed that the MOSPD2 mutant devoid of transmembrane domain (TM) remained cytosolic and did not accumulate on the LD surface (Fig. 5B and C). To better understand this, TM domain of MOSPD2 was substituted by the TM of the ER-anchored phosphatase SAC1 that comprises two transmembrane helices (named TM(SAC1)) (Fig. 5E). While the fusion protein GFP-TM(SAC1) was evenly localized in the ER, the chimeric protein composed of MOSPD2 CRAL-TRIO and MSP domains fused with the TM(SAC1) domain was present in the ER and accumulated around LDs. Similarly, the fusion protein comprising only the CRAL-TRIO domain of MOSPD2 and the TM(SAC1) domain was also present in the ER and enriched around LDs (Fig. 5E). Thus, MOSPD2 needs an ER-anchor to be localized in contact with LDs.

Together, these data show that the TM and CRAL-TRIO domains are necessary for MOSPD2 binding to LDs.

### An amphipathic helix in the CRAL-TRIO domain of MOSPD2 is required for binding to LDs

Proteins that associate with LDs do so via two modalities at least: Class I proteins are embedded in the ER bilayer and can diffuse laterally to the LD monolayer, while Class II proteins translocate from the cytosol to the surface of LDs (Olzmann & Carvalho, 2019; Kory *et al*, 2016). Most Class II proteins associate with LDs through an amphipathic α-helix (AH), in which hydrophobic and polar residues are segregated to form two distinct faces along the helix axis. This topology allows AH to efficiently bind membranes because hydrophobic residues can insert between lipid acyl chains whereas polar residues can make polar contacts with lipid headgroups. The LD surface has more packing defects than a bilayer, i.e. gaps in the phospholipid layer, which are favorable for the insertion of hydrophobic residues and thus the association of Class II proteins (Giménez-Andrés *et al*, 2018; Chorlay & Thiam, 2020; Chorlay *et al*, 2021). Because MOSPD2 is anchored to the ER and does not diffuse to the LD monolayer, we hypothesized that the CRAL-TRIO domain might behave like a Class II protein and thus possess an AH. As no experimental structure of the CRAL-TRIO domain of MOSPD2 was available, we built structural models of the protein using SWISS-MODEL and AlphaFold (Waterhouse *et al*, 2018; Jumper *et al*, 2021), and identified α helices in the models. We determined their hydrophobicity and hydrophobic moment using Heliquest (Gautier *et al*, 2008) and identified an AH at the end of the CRAL-TRIO domain (Fig. 6A and B, and Fig. S8A), exposed at the surface of the protein and thus potentially able to insert into a membrane. Sequence analyses showed that the helix is highly conserved from Cnidaria to Human (Fig. 6C). To determine whether this AH is responsible for MOSPD2 binding to LDs, we replaced the bulky hydrophobic residue tryptophan 201 in the middle of the nonpolar face by a negatively charged residue glutamate (mutant W201E) which would perturb the membrane partitioning of this helix. Compared with WT-MOSPD2 which is found both in the ER and in ER-LD contacts when expressed in cells, MOSPD2 W201E mutant was evenly distributed in the ER (Fig. 6D and E). Moreover, replacing the CRAL-TRIO domain by the AH only (GFP-AH-MOSPD2 ΔCRAL-TRIO fusion protein) was sufficient to recruit MOSPD2 on LDs (Fig. 6D and E).

**Figure 6:**
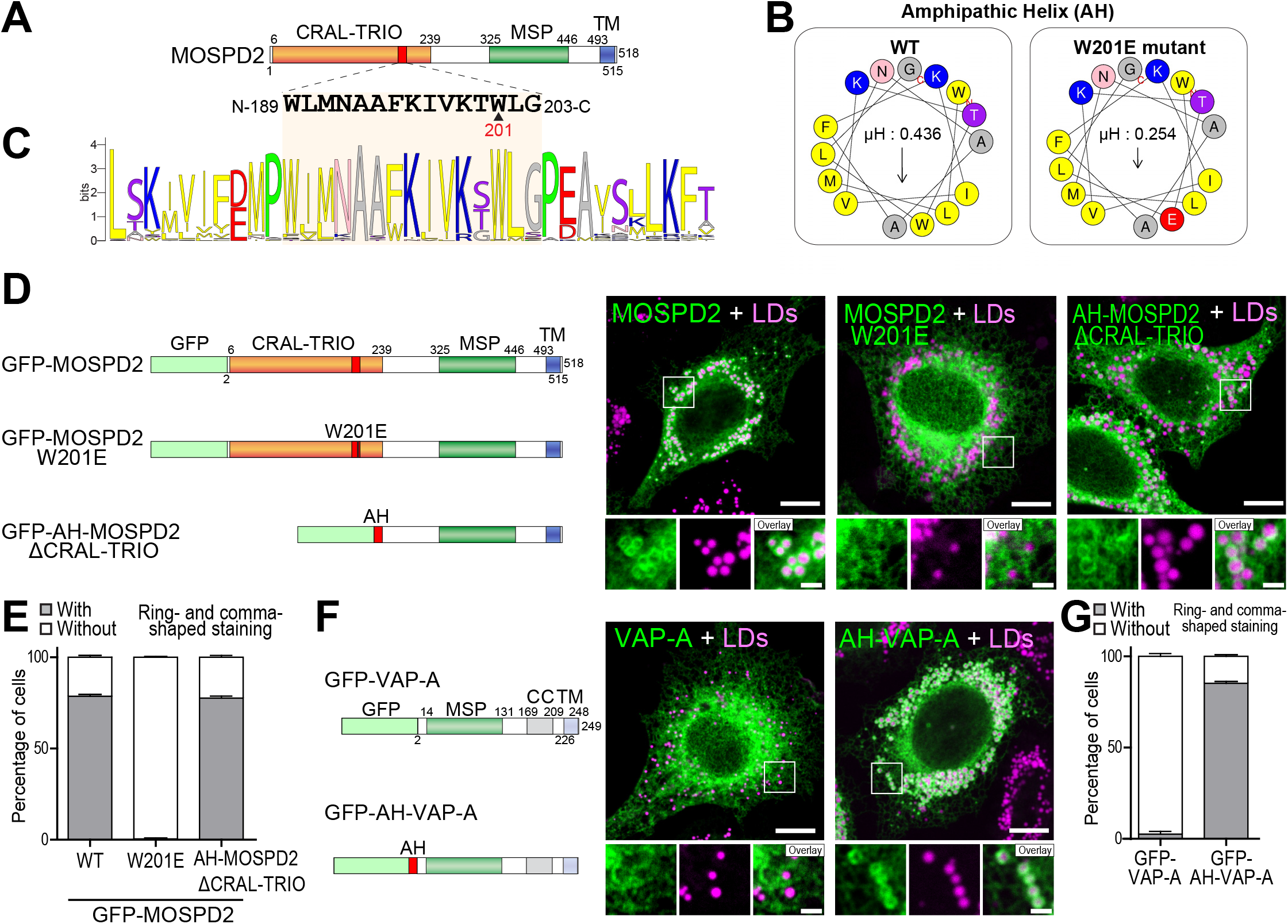
An amphipathic helix in the CRAL-TRIO domain of MOSPD2 mediates its localization at ER-LD contacts. A. Schematic representation of MOSPD2 showing the position of the amphipathic helix (red) and its sequence. The arrowhead shows the position of residue W201. B. Helical wheel representation of the WT (left) and W201E mutant (right) AH (aa 189-203) generated with HeliQuest (http://he-liquest.ipmc.cnrs.fr/) (left part). The W201E mutation alters the amphipathic character of the helix by reducing its hydrophobic moment (μH) from 0.436 to 0.254. C. WebLogo generated from an alignment of MOSPD2 AH sequence from 44 species. The AH is highlighted in light orange and 10 flanking residues from either side are shown. D. Left: schematic representation of GFP-MOSPD2 constructs either WT (GFP-MOSPD2) or bearing a mutation in the AH (GFP-MOSPD2 W201E), or containing a deletion of the CRAL-TRIO domain together with an insertion of the AH (AH-MOSPD2-Δ CRAL-TRIO). Right: localization of these constructs in HeLa cells treated with OA; LDs were stained with Nile Red (magenta). Images were acquired on a confocal microscope (Leica SP5; 63x NA 1.4). Scale bars: 10 μm (insets 2 μm). E. Quantification of cells showing ring- or comma-shaped staining for these constructs. Mean ± SD; n=3 independent experiments (MOSPD2 WT:117 cells; W201E: 156 cells; AH-MSP-TM: 113 cells). F. Localization of WT GFP-VAP-A and GFP-AH-VAP-A chimera in which the AH of MOSPD2 was fused at the N-terminus of VAP-A. LDs were stained with Nile Red in HeLa cells treated with OA. Confocal microscope (Leica SP5; 63x NA 1.4) images. Scale bars: 10 μm (insets 2 μm). G. Quantification of signal localization in cells expressing WT GFP-VAP-A and GFP-AH-VAP-A chimera showing ring- or comma-shaped staining. Mean ± SD; n = 3 independent experiments (VAP-A WT: 109 cells; AH-VAP-A: 102 cells). Data information: In (D, F), composite subpanels on the bottom are higher magnification images of the area outlined. The overlay panel shows merged channels.

As mentioned before, VAP-A is not recruited in ER-LD contacts (Fig. 1B and Fig. 4E). To know whether the AH of MOSPD2 could allow the recruitment of VAP-A on LDs, we created a chimeric protein composed of the AH of MOSPD2 fused to VAP-A. Unlike VAP-A, which is distributed evenly in the ER, the fusion protein AH-VAP-A accumulated in ER subdomains around LDs (Fig. 6F and G).

Combined together, these data show that the AH of MOSPD2 is necessary and sufficient for this ER-bound protein to mediate the formation of ER-LD contacts.

### The amphipathic helix of MOSPD2 directly interacts with the surface of LDs

To directly test whether the AH of MOSPD2 could bind LDs, we carried out flotation assays using artificial LDs (aLDs) and fluorescein isothiocyanate (FITC)-labeled synthetic peptides encompassing the AH of MOSPD2, either wild type or with the W201E mutation (Fig. 7A). A peptide with a random sequence was used as negative control (Fig. 7A). aLDs composed of a mix of triolein and surrounded by a monolayer of POPC and POPE labeled with a fluorescent lipid (Rhodamine-PE) were prepared. The peptides were incubated with these aLDs, mixed with sucrose, and allowed to float over this sucrose cushion (Fig. 7B). Three fractions corresponding to the top, middle and bottom position of the cushion were collected, and the fluorescence signal of aLDs and of the peptides were measured. After ultracentrifugation, Rhodamine-labelled aLDs were in the top fraction (Fig. 7C). The control peptide remained in the bottom fraction, while the peptide corresponding to the AH of MOSPD2 was in the top fraction with aLDs. Unlike the WT AH of MOSPD2, the W201E mutant behaved like the negative control peptide and remained in the bottom fraction. To further characterize the association of the AH of MOSPD2 to aLDs, we performed aLD - peptide interaction assays. aLDs composed of triolein were mixed with the fluorescent peptides. aLDs were imaged and fluorescence on the surface of aLDs quantified. In agreement with the flotation assays, the control peptide did not bind to the aLDs, while the peptide corresponding to the AH of MOSPD2 was found attached to the aLDs. The W201E mutant was only minimally retained by the aLDs. Together, these experiments show that the AH of MOSPD2 directly binds to LDs.

**Figure 7:**
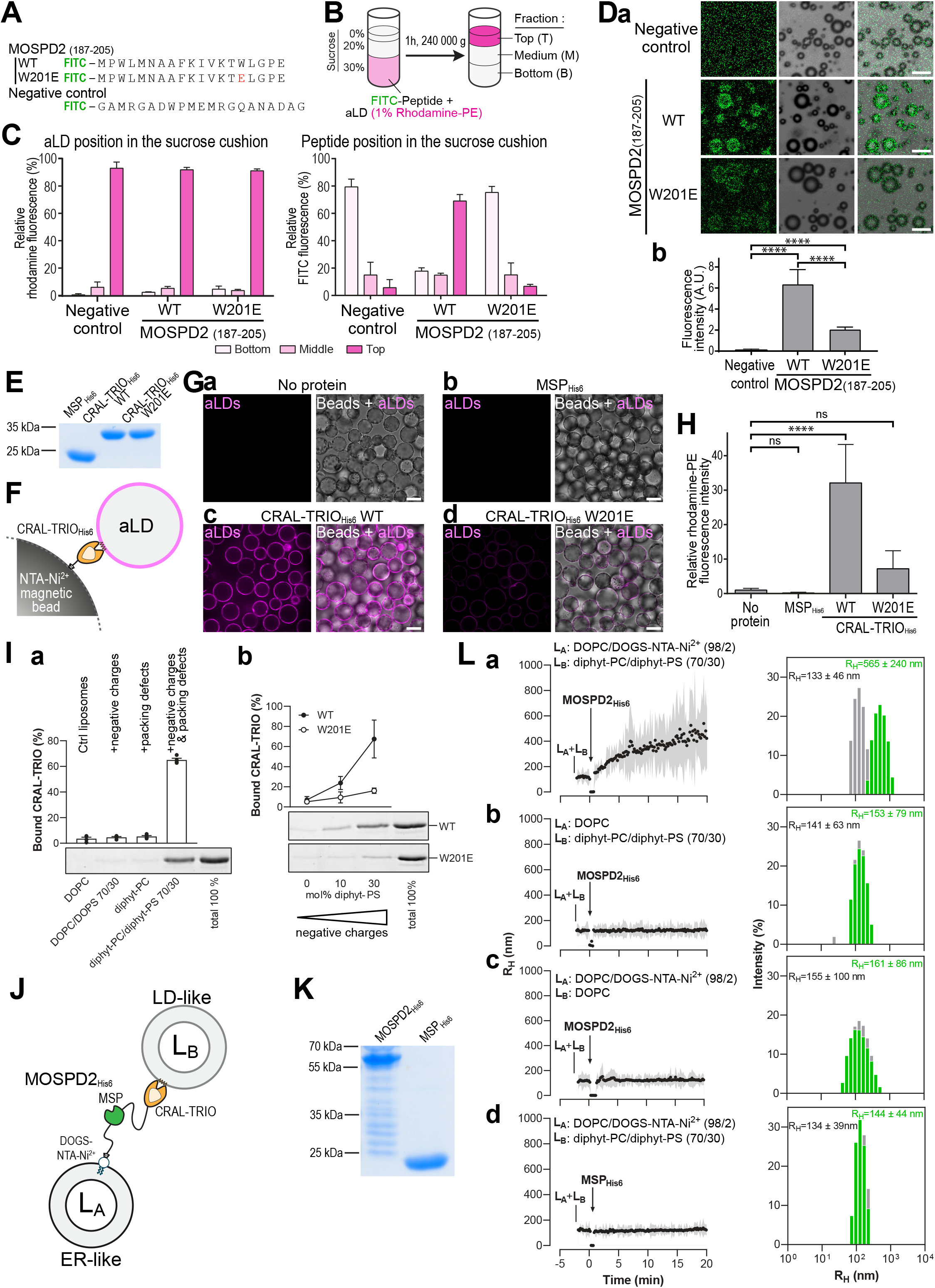
The CRAL-TRIO domain of MOSPD2 directly interacts with aLDs. A. Peptides used for artificial LDs (aLDs) flotation assays. Peptides corresponding to the WT or W201E mutant AH of MOSPD2 (residues 187-205), and negative control composed of a random sequence, were coupled with Fluorescein (FITC) at their amino-terminal end. B. Principle of artificial LDs (aLDs) flotation assays. Fluorescent peptides were incubated with aLDs containing 1 mol% Rhodamine-PE, then ultracentrifuged to allow aLDs to float on the sucrose cushion. Top, middle and bottom fractions were collected and FITC and rhodamine fluorescence quantified. C. aLDs flotation assays. Left: relative rhodamine fluorescence (*i.e*. aLDs); right: relative FITC fluorescence (*i.e*. peptides), in the bottom (light pink), middle (pink) and top (dark pink) fractions. Means (± SD) from n = 5 independent experiments. D. aLDs - peptide interaction assay. a: Representative images of aLDs incubated with peptides shown in A. Scale bars: 10 μm. b: quantification of peptide fluorescence on aLDs (Means (±SD); negative control: 1397, WT AH of MOSPD2: 231, W201E mutant AH of MOSPD2 198 aLDs from 2 independent experiments). Student’s t-test (****: P<0.0001). E. Coomassie blue staining of the recombinant MSP_His6_, WT CRAL-TRIO_His6_ and mutant CRAL-TRIO_His6_ W201E proteins after SDS-PAGE. F. Principle of aLDs pull-down assay. Proteins were immobilized on magnetic NTA-Ni^2+^ beads, owing to their His6 tag, and incubated with fluorescent aLDs. G. Representative confocal images of NTA-Ni^2+^ beads either not coated with recombinant proteins (a, no protein), and coated with recombinant domains of MOSPD2 (b, MSP_His6_; c, WT CRAL-TRIO_His6_; and d, mutant CRAL-TRIO_His6_ W201E) and incubated with fluorescent aLDs (magenta). Left: confocal section of aLD fluorescence; right: superposition with brightfield images showing the beads. Spinning disk confocal microscope (Nikon CSU-X1, 100x NA 1.4) images. Scale bars: 10 μm. H. Quantification of aLDs recruitment on NTA-Ni^2+^ beads. Rhodamine fluorescence was measured using a fluorimeter. Means (±SD). Kruskal-Wallis with Tukey’s multiple comparisons test (****: P<0.0001; ns: not significant; n = 6 independent experiments). I. a: Flotation assays. CRAL-TRIO_His6_ (0.75 μM) was mixed with liposomes (0.75 mM lipids) only made of DOPC or diphyt-PC or composed of DOPC/DOPS (7/3 mol/mol) or diphyt-PC/ diphyt-PS (7/3 mol/mol) in HK buffer at 25°C for 10 minutes. After centrifugation, the liposomes were recovered at the top of a sucrose cushion and analyzed by SDS-PAGE. The amount of protein recovered in the top fraction (lane 1 to 4) was quantified and the fraction of liposome-bound CRAL-TRIO_His6_ was determined using the content of lane 5 (total 100%) as a reference. Data are represented as mean ± SEM (error bars; n = 4). b: Flotation assays. WT (closed circle) and W201E mutant (open circle) MOSPD2 CRAL-TRIO_His6_ proteins (0.75 μM) were mixed for 10 minutes with liposomes (0.75 mM lipids) only made of diphyt-PC or additionally containing 10 or 30 mol% diphyt-PS. Data are represented as mean ± SEM (error bars; n = 3-5) J. Principle of the membrane tethering assay. K. Coomassie blue staining of the recombinant MOSPD2_His6_ and MSP_His6_ proteins after SDS-PAGE. L. Membrane tethering assays. L_A_ liposomes (50 μM total lipids) composed of DOPC/DOGS-NTA-Ni^2+^ (98/2 mol/mol) (a, b and d) or DOPC (b) were mixed with LB liposomes (50 μM), composed of diphyt-PC/diphyt-PS (70/30 mol/mol) (a, b and d) or DOPC (c) in HK buffer at 25°C. After 2 minutes, MOSPD2_His6_ (a, b and c) or MSP_His6_ (d) (0.4 μM) was added and the size of liposomes was measured for 23 minutes. Left panels: Mean radius (dots) and polydispersity (shaded area) over time. Right panels: R_H_ distribution before (gray bars) and after the reaction (green bars). These experiments are representative of several independent experiments (n = 3-5).

Then, we tested whether the CRAL-TRIO domain binds to LDs via its AH. We produced in *Escherichia coli* and purified the wild type and W201E mutant CRAL-TRIO domains of MOSPD2 fused with a His6 tag (Fig 7D). We also purified the MSP domain of MOSPD2 fused with a His6 tag as a control (Fig. 7D). By circular dichroism (CD) spectroscopy, we established that the content in secondary structure of the W201E mutant was identical to that of the wild-type WT CRAL-TRIO domain, indicating that the mutation did not impair the folding of the domain (Fig. S8B). To assess the ability of these three recombinant proteins to bind aLDs, we performed aLD pull-down assays. Each protein was immobilized on magnetic NTA-Ni^2+^ beads, owing to its His6 tag, and incubated with fluorescent aLDs (Fig. 7E). After several washes to remove unbound aLDs, the beads were imaged (Fig. 7F) and fluorescence quantified using a fluorimeter (Fig. 7G). In the absence of protein or in the presence of the MSP_His6_, no fluorescence was measured, meaning that aLDs were not retained on the beads (Fig. 7F, a and b, and 7G). In contrast, when the wild type CRAL-TRIO_His6_ was attached to the beads, a high fluorescence was detected showing that aLDs were retained by the protein (Fig. 7F, c and 7G). In comparison, a much lower aLD retention was observed with the W201E mutant (Fig. 7F, d and 7G). These data indicate that the AH of the CRAL-TRIO domain is instrumental for the protein to bind LDs.

To better define which membrane determinants facilitate MOSPD2 binding to LDs, we performed flotation assays with membranes that differ in terms of lipid packing defect and electrostatics (Fig. 7H). The recombinant CRAL-TRIO domain of the protein was tested with different types of liposomes made of phospholipids and with or without negative charges and/or packaging defects. Control liposomes with few packing defects and no charge were composed of phosphatidycholine with di-oleyl (DOPC). Negative charges were provided by replacing 30 % of phosphatidylcholine by the anionic lipid phosphatidylserine (PS). Finally, packing defects were generated by using phospholipids containing diphytanoyl (diphyt-PC) acyl chains; diphytanoyl is a 16:0 acyl-chain with branched methyl groups that forms large packing defects. The CRAL-TRIO_His6_ protein was poorly bound by control liposomes (no charges, no packing defects). It did not associate either with liposomes having negative charges only (DOPC/DOPS 70/30), or having packing defects only (diphyt-PC) (< 6 % membrane-bound protein) (Fig. 7H, a). In contrast, more than 60% of the protein was associated with liposomes containing both negative charges and packing defects (diphyt-PC/diphyt-PS 70/30) (Fig. 7H, a). Flotation assays performed with liposomes bearing packing defects and increasing concentration of negatively charged phospholipids showed that the binding of the CRAL-TRIO_His6_ protein was proportional to the amount of charges (Fig. 7H, b). Thus, the binding of the CRAL-TRIO domain to packing defects bearing liposomes is tuned by electrostatics. Noteworthy, almost no binding was seen with MOSPD2 CRAL-TRIO W201E even in the presence of more packing defects and negative charges (Fig. 7H, b). Collectively these data show that in vitro the association of the CRAL-TRIO domain of MOSDP2 with a membrane is facilitated by the presence of very large packing defects and negatively charged lipids, both of which are characteristics of the LD surface.

Finally, we examined *in vitro* whether MOSPD2 was able to directly connect the ER with LDs by performing membrane tethering assays. These were done using two populations of liposomes, L_A_ and L_B_, mimicking the ER and LDs, respectively. The association of liposomes into large particles as a result of membrane tethering was measured by dynamic light scattering (DLS). L_A_ liposomes made of DOPC and doped with DOGS-NTA-Ni^2+^ were mixed with L_B_ liposomes composed of diphyt-PC/diphyt-PS (70/30 mol/mol) (Fig. 7I). Then, MOSPD2_His6_, corresponding to the cytosolic part of MOSPD2 tagged with a C-terminal 6His-tag (Fig. 7J), was added so that L_A_ liposomes were covered by the protein and constituted ER-like liposomes. A rapid increase in the initial mean radius of liposomes was observed suggesting that MOSPD2, once attached to L_A_ liposomes, connected them with L_B_ liposomes (Fig. 7K, a). In contrast, no aggregation occurred when L_A_ liposomes were devoid of attached MOSPD2_His6_ (Fig. 7K, b), or when they were covered by the MSP domain of MOSPD2 (MSP_His6_) (Fig. 7K, c). Moreover, no aggregation was observed when L_B_ liposomes were replaced with liposomes that did not mimic LDs (i.e. without packing defects and negative charges) (Fig. 7K, d).

These data showed that MOSPD2, anchored to the ER by its C-terminus, directly connects this compartment with a second one delimited by a membrane with large packing defects and anionic lipids, such as LDs, owing to its CRAL-TRIO domain.

### The formation of ER-LD contacts mediated by the CRAL-TRIO domain is essential for the function of MOSPD2 in the biology of LDs, while the MSP domain is dispensable

In the absence of MOSPD2, lipid droplets are fewer (Fig. 2C). Having identified the molecular mechanism of ER-LD contact formation mediated by MOSPD2, we asked whether its ability to form ER-LD contacts was required for its role in LD biology. To answer this question, we performed rescue experiments by restoring MOSPD2 expression in knock-out cells (KO#1) using a mScarlet-tagged WT or mutant MOSPD2 (Fig. 8 A, B). LD were then labeled and their number and size quantified (Fig. 8 C, D). Consistent with data from Fig. 2B and C, MOSPD2 knock-out cells had two times fewer LDs than wild-type cells (Fig. 8C, a-b and 8D). When mScarlet-MOSPD2 was re-expressed in knock-out cells, the number of LDs was similar to that of WT cells (Fig. 8C, c and 8D). Thus, the ectopic expression of MOSPD2 rescues the LD phenotype of MOSPD2 knock-out cells.

**Figure 8:**
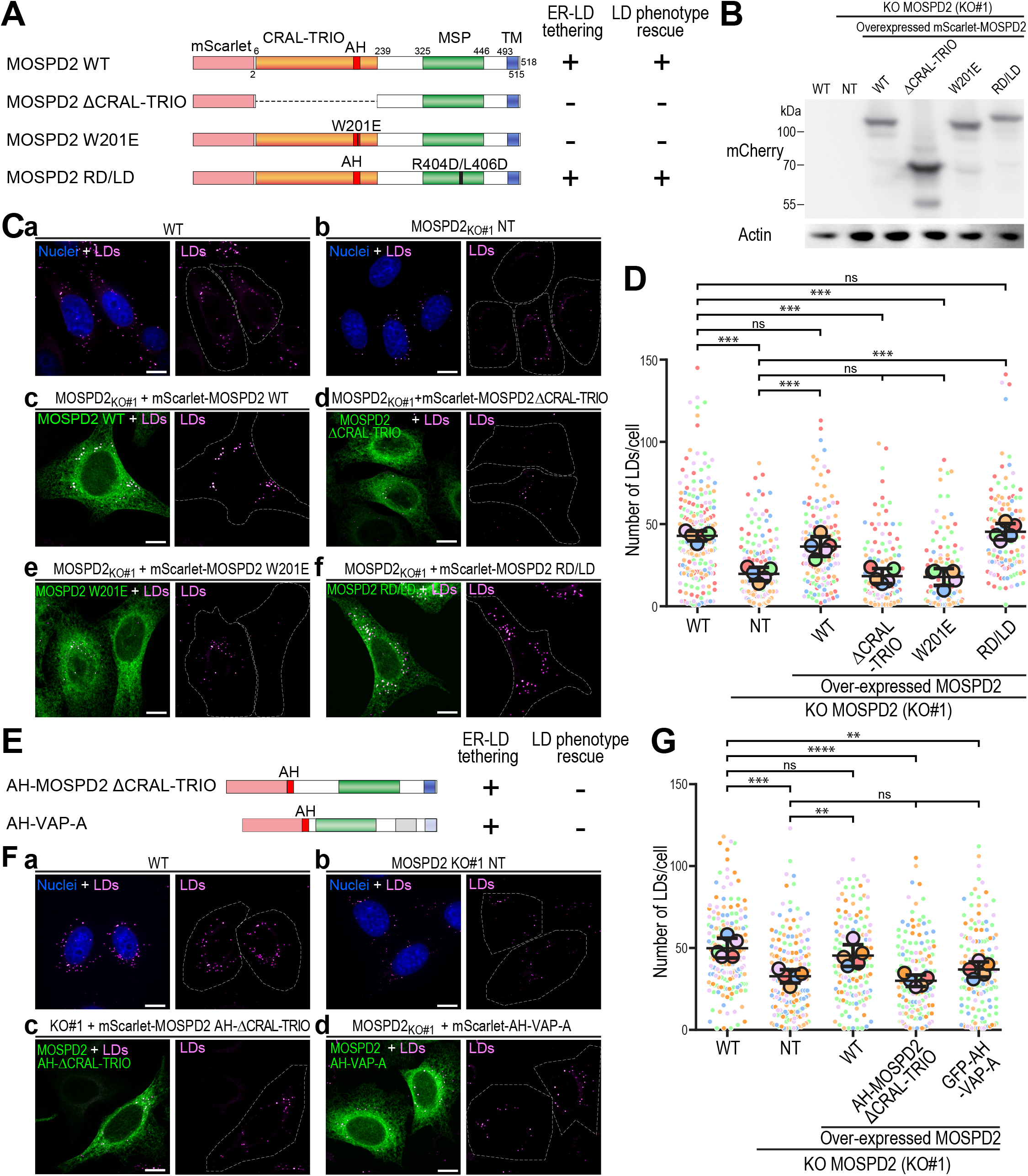
The capacity of MOSPD2 to form ER-LD contact sites is necessary but not sufficient to regulate LDs. A. Schematic representation of mScarlet-MOSPD2 constructs either WT (mScarlet-MOSPD2) or containing a deletion of the MSP domain (Δ MSP) or the CRAL-TRIO domain (ΔCRAL-TRIO), a mutation in the MSP domain (RD/LD) or in the CRAL-TRIO domain (W201E). For each constructs, the LD tethering activity and the rescue (see panels below) are summarized as + or -. B. Western Blot analysis of WT and MOSPD2 knock-out (KO#1) HeLa cells. MOSPD2 expression was rescued in MOSPD2 knock-out cells using mScarlet-MOSPD2 expression constructs either WT or mutant (mScarlet-MOSPD2 ΔCRAL-TRIO, W201E and RD/LD). NT: non-transfected. C. Representative confocal images of WT and MOSPD2 knock-out (KO#1) HeLa cells in which MOSPD2 expression was restored using mScarlet-MOSPD2 constructs (green) depicted in panel A. As control, untransfected WT (a) and MOSPD2 knock-out (b) cells were imaged. LDs were stained with BODIPY 493/503 (magenta) and nuclei with Hoechst (blue). Images were acquired on a spinning disk confocal microscope (Nikon CSU-X1, 100x NA 1.4). Scale bars: 10 μm. D. Quantification of the number of LDs in cells shown in B. Data are displayed as Superplots showing the mean number of LDs per cell (small dots), and the mean number of LDs per independent experiment (large dots). Independent experiments (n = 5) are color-coded. Means and error bars (SD) are shown as black bars. Data were collected from 213 (WT), 200 (KO#1), 150 (KO+mScarlet-MOSPD2 WT), 238 (KO+mScarlet-MOSPD2 ΔCRAL-TRIO), 126 (KO+mScarlet-MOSPD2 W201E), and 118 (KO+mScarlet-MOSPD2 RD/LD) cells. One-way ANOVA with Tukey’s multiple comparisons test (***: P < 0.001; ns: not significant; n = 5 independent experiments). E. Schematic representation of mScarlet constructs containing the deletion of the CRAL-TRIO domain together with an insertion of the AH (AH-MOSPD2-ΔCRAL-TRIO), and of the GFP-AH-VAP-A chimera in which the AH of MOSPD2 was fused at the N-terminus of VAP-A. For both constructs, the LD tethering activity and the rescue (see panels below) are summarized as + or -. F. Representative confocal images of WT and MOSPD2 knock-out (KO#1) of HeLa cells in which constructs (green) from panel E were expressed. As control, untransfected WT (a) and MOSPD2 knock-out (b) cells were imaged. LDs were stained with BODIPY 493/503 (magenta) and nuclei with Hoechst (blue). Images were acquired on a spinning disk confocal microscope (Nikon CSU-X1, 100x NA 1.4). Scale bars: 10 μm. G. Quantification of the number of LDs in cells shown in F. Data are displayed as Superplots showing the mean number of LDs per cell (small dots), and the mean number of LDs per independent experiment (large dots). Independent experiments (n = 5) are color-coded. Means and error bars (SD) are shown as black bars. Data were collected from 202 (WT), 192 (KO#1), 156 (KO+mScarlet-MOSPD2 WT), 147 (KO+mScarlet-MOSPD2 AH-ΔCRAL-TRIO) and 155 (KO+mScarlet-AH-VAP-A). One-way ANOVA with Tukey’s multiple comparisons test (*: P<0.05; ***: P < 0.001; ns: not significant; n = 5 independent experiments).

Next, we repeated the rescue experiment by expressing two MOSPD2 mutants unable to form ER-LD contacts, a deletion mutant devoid of CRAL-TRIO domain (mScarlet-MOSPD2 ΔCRAL-TRIO) and a point mutant with a defective AH (mScarlet-MOSPD2 W201E), in MOSPD2 knock-out cells (Fig. 8A, B). These two mutants failed to rescue the absence of MOSPD2 (Fig. 8Ce-f, and 8D).

In contrast, the expression of a mutant MOSPD2 having an MSP domain unable to bind FFAT motifs (mScarlet-MOSPD2 RD/LD) restored the number of LDs to a level similar to that of WT cells (Fig. 8Cf and 8D). Thus, the ability of MOSPD2 to bind FFAT motifs is dispensable for its function in LDs. These experiments show that the ability of MOSPD2 to form ER-LD contacts by its CRAL-TRIO domain contributes to LD biology.

Finally, we tested whether promoting ER-LD tethering was sufficient to recapitulate the function of MOSPD2. We expressed two constructs lacking a CRAL-TRIO domain but capable of promoting ER-LD contact formation (Fig. 6): a MOSPD2 mutant in which the CRAL-TRIO domain was replaced by the AH only (GFP-AH-MOSPD2 ΔCRAL-TRIO fusion protein), and the chimeric protein composed of the AH of MOSPD2 fused to VAP-A (Fig. 8F). While these two proteins promoted ER-LD contact formation (Fig. 6 D-G), they did not rescue the number of LDs in MOSPD2-deficient cells (Fig. 8F, G). These data show that MOSPD2 must connect the ER with the LDs to exert its activity, but that the sole membrane tethering ability is not sufficient to recapitulate the activity of MOSPD2.

Together, these data show that the ability of MOSPD2 to create ER-LD contacts is crucial for the activity of the protein in the biology of LDs. Moreover, it shows that the CRAL-TRIO domain is instrumental for MOSPD2 function in LDs, while the MSP domain is dispensable.

## Discussion

Organelles are no longer considered as isolated compartments but as active units able to constantly communicate and function with each other. The ER plays a key role in the interactions between organelles, as it is a meshwork of membrane tubes and sheets that extent throughout the cytosol and make extensive contacts with other organelles (Wu *et al*, 2018). LDs, which are cellular energy storage, have a very unique relationship with the ER: they are generated from the ER and maintain regular physical contacts with it throughout their life cycle (Olzmann & Carvalho, 2019; Walther *et al*, 2017). LD biogenesis starts with the synthesis of neutral lipids in the ER membrane. These newly made lipids nucleate into oil lenses in the ER bilayer that ultimately bud towards the cytosol and further grow. At the beginning of their life, LDs are attached to the ER as their monolayer is continuous with the cytosolic leaflet of the ER membrane (Hugenroth & Bohnert, 2020; Salo & Ikonen, 2019). Eventually, LDs bud towards the cytosol and detach from the ER but continue to establish contacts with the ER by other mechanisms. This physical connectivity might ensure the functional interplay between the ER and LDs. In this study, we reveal that MOSPD2 contribute to this process by forming contacts between the ER and LDs, and participates to the biology of these organelles (Fig. 9).

**Figure 9:**
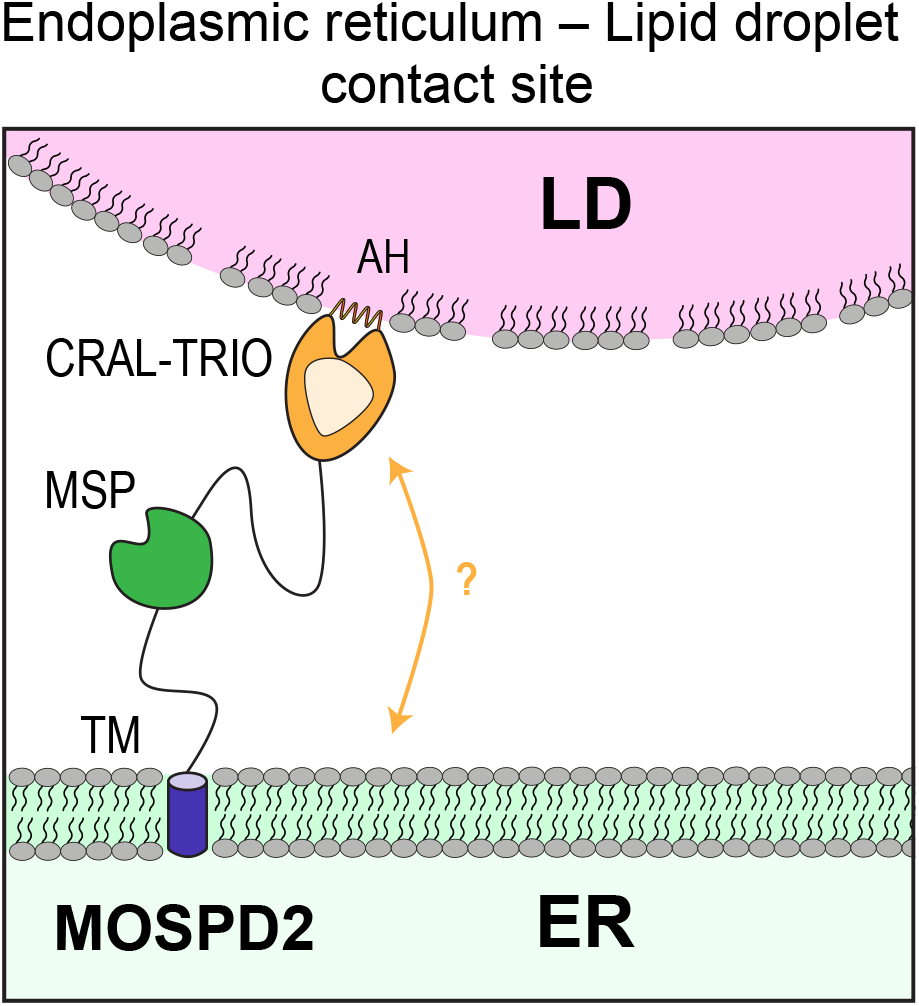
Schematic representation of ER-LD contact sites mediated by MOSPD2. MOSPD2 tethers the ER to LDs thanks to its TM and CRAL-TRIO domains. The amphipathic helix located in the CRAL-TRIO domain directly interacts with the surface of LDs. The CRAL-TRIO domain of MOSPD2 might also be involved in lipid transport between the ER and LDs.

We previously showed that MOSDP2 mediates the formation of contacts between the ER and endosomes, the Golgi and mitochondria (Di Mattia *et al*, 2018) by a mechanism relying on its MSP domain. By binding to FFAT motifs present in proteins on the surface of these organelles, the MSP domain of MOSPD2 attaches the ER to the other organelle, as do VAP-A and VAP-B proteins (Di Mattia *et al*, 2018, 2020a; Murphy & Levine, 2016). Here, we identified that MOSPD2 has a second tethering activity to specifically create ER-LD contacts by a mechanism that does not rely on its MSP but surprisingly on its CRAL-TRIO domain.

The association of this domain with LDs is mediated by an amphipathic helix that is conserved in other members of the CRAL-TRIO family (Bankaitis *et al*, 2010). and notably its archetypical member, Sec14p (Sha *et al*, 1998). In this protein, this helix acts as a gate regulating access to the hydrophobic cavity of the protein (Ryan *et al*, 2007; Bankaitis *et al*, 1990; Sha *et al*, 1998). AHs are found in a variety of proteins interacting with membranes; they do not have a consensus sequence, and they were shown to interact with diverse polar/non-polar interfaces (Giménez-Andrés *et al*, 2018). Depending on the type of hydrophobic residues on their apolar side, the nature and charge of residues on their polar side and their length, they can interact with membranes having distinct properties in terms of lipid composition, charge, curvature… (Giménez-Andrés *et al*, 2018). AHs binding to LDs have affinity for packing defects owing to the presence of large hydrophobic residues on their apolar face (Prévost *et al*, 2018; Chorlay & Thiam, 2020; Čopič *et al*, 2018). In agreement with this notion, mutation of W201 in the AH of MOSPD2 abolishes its binding to LDs. Besides, the CRAL-TRIO domain of MOSDP2 has no or little affinity for other cellular membranes such as the ER membrane which is expected to be more packed than the LD surface and contain no more than 15 mol% negatively-charged lipids (van Meer *et al*, 2008; Chorlay & Thiam, 2020; Chorlay *et al*, 2021).

The main feature of CRAL-TRIO domain is to possess a cavity to specifically host lipophilic molecule. CRAL-TRIO domain containing proteins belong to the large category of lipid transfer proteins (Chiapparino *et al*, 2016). Sec14 can exchange glycerophospholipids (PI and PC) between membranes, whereas retinaldehyde-binding protein 1 (RLBP1) and alpha-tocopherol transfer protein (TTPA) use a CRAL/TRIO domain to convey retinaldehyde and α-tocopherol (vitamin E), respectively. Thus, an appealing hypothesis is that MOSPD2 transports fatty acid to fuel LD enzymes that build neutral lipids, or transport phospholipids to the monolayer of LDs to allow their proper growth. The reduced level of sterol esters in MOSPD2-deficient cells points to a link between MOSPD2 and the metabolism of cholesterol and its derivatives. Therefore, one possibility is that MOSPD2 could transport sterol towards LDs. Interestingly, other lipid transfer proteins are present in ER-LD contacts such as ORP5 and ORP8 (Guyard *et al*, 2021; Du *et al*, 2019), therefore raising the possibility that ER-LD contacts are a major platform for the non-vesicular exchange of lipids. If MOSPD2 is a lipid transporter, we can speculate that the AH of the CRAL-TRIO domain of MOSPD2 has a dual function: interacting with the surface of LDs to mediate the formation of ER-LD contacts, and being a gate to access the hydrophobic cavity of the CRAL-TRIO domain of the protein. However, a debate still exists on whether Sec14p exchanges lipids between organelles in yeast. As proposed for Sec14p, rather than transporting lipids, MOSPD2 could act by presenting its lipid substrate to an enzyme and thus increase its activity (Lete *et al*, 2020). Further work will be needed to determine if MOSPD2 mediates lipid transport at the ER/LD interface.

It is intriguing that a single protein possesses two tethering mechanisms targeting distinct organelles: on the one hand, the MSP domain which allows the formation of contacts with many organelles via protein-protein interactions, and on the other hand the CRAL-TRIO domain which contacts only one organelle, the LD, via a protein-membrane interaction. The ability of MOSPD2 to tether the ER to other organelles using two alternate molecular mechanisms is a new illustration of the plasticity of inter-organelle contacts. Indeed, other tether proteins alternate between distinct contacts. For instance, the mitochondria-bound protein MIGA2 is present in mitochondria-ER contacts when bound to VAP proteins thanks to its Phospho-FFAT motif, and alternatively in mitochondria-LD contacts by directly binding to the surface of LDs thanks to an amphipathic helix. These two localizations of MIGA2 most probably reflect alternate functions of the protein related to cellular metabolism (Klemm, 2021; Freyre *et al*, 2019). Similarly, some members of the VPS13 family are present in different inter-organelle contacts, in contacts with LDs using an AH (Yeshaw *et al*, 2019; Wang *et al*, 2021; Kumar *et al*, 2018; Ramseyer *et al*, 2018), and with the ER using FFAT and Phospho-FFAT (Guillén-Samander *et al*, 2021; Wang *et al*, 2021). In the case of MOSPD2, the deletion of the MSP domain, or its mutation rendering it unable to bind to FFAT motifs, unexpectedly promoted the formation of ER-LD contacts. This observation suggests that MOSPD2 is balanced between two kinds of membrane contact sites: MSP domain-dependent contacts that involve FFAT-containing partners of MOSPD2, and CRAL-TRIO domain-dependent contacts that involve the direct recognition of the surface of LDs by MOSPD2. It is plausible that regulation mechanisms exist to control the repartition of MOSPD2 between the two types of contact sites in which the protein might then play distinct roles. In line with this, we have recently shown that phosphorylation of FFAT-like motifs that we named Phospho-FFAT allows a regulatable MSP-dependent formation of contact sites (Di Mattia *et al*, 2020a). Thus, the Phospho-FFAT phosphorylation status of MOSPD2 partners can probably indirectly control pools of MOSPD2 in and out ER-LD contacts. We also observed that in cells producing large amounts of LDs, MOSPD2 accumulated in ER-LD contacts, suggesting that ER-LD contacts are privileged, and that the metabolic state of the cell probably dictates the localization and function of MOSPD2. Moreover, the binding of MOSPD2 to LDs might also be tuned by the lipid composition of LDs; in agreement with our in vitro data, changes in the phospholipid monolayer or in the neutral core of LDs most probably regulate the affinity of binding of MOSPD2. However, it is unclear whether the association of MOSPD2 with LDs is at the expense of other specific contacts or simply reduces the amount of MOSPD2 that can be recruited by FFAT-containing partners. In either case, this would represent an additional level of control of contacts between the ER and other organelles.

To summarize, we report that MOSPD2 builds ER-LD contacts and thereby affects LD homeostasis. MOSPD2 shares many similarities with VAP-A and VAP-B: these three proteins recognize FFAT and Phospho-FFAT motifs, they have many partners in common, and by this molecular mechanism are involved in the formation of contacts between the ER and many organelles. The finding that MOSPD2 makes additional contacts through a mechanism distinct to that of VAP-A and VAP-B provides specificity and expands the repertoire of contacts that this protein can make. The formation of contacts between the ER and other organelles is a plastic phenomenon involving complex networks of interactions, and future work should provide evidence on how the recruitment of MOSPD2 in a given inter-organelle contact is orchestrated.

## Materials and Methods

### Cloning and constructs

The GFP-MOSPD2 (WT and RD/LD mutant), GFP-VAP-A and GFP-TM(SAC1) expression vectors were previously described (Alpy *et al*, 2005, 2013; Di Mattia *et al*, 2018).

The GFP-MOSPD2 ΔMSP, GFP-MOSPD2 ΔTM, GFP-MOSPD2 ΔCRAL-TRIO, GFP-MOSPD2 W201E expression vectors were constructed by overlap extension polymerase chain reaction (PCR) using GFP-MOSPD2 as a template and the following central primers: GFP-MOSPD2 ΔMSP: 5’-AGT GTA TTT AAA GGC CCC GAA AGC AGT AAA CCA AAC-3’ and 5’-GTT TGG TTT ACT GCT TTC GGG GCC TTT AAA TAC ACT-3’; GFP-MOSPD2 ΔTM: 5’-CAG CGT TGT ATC TGA ATT CCA GCA GCT GCT GCT TTC C-3’ and 5’-CAG CTG CTG GAA TTC AGA TAC AAC GCT GAA CTT GGT C-3’; GFP-MOSPD2 ΔCRAL-TRIO: 5’-ATC ATC TAC TAG TGG TGG ATA GCT AGA ATT CGA AGC TTG AGC TCG AGA-3’ and 5’-TCT CGA GCT CAA GCT TCG AAT TCT AGC TAT CCA CCA CTA GTA GAT GAT-3’; GFP-MOSPD2 W201E: 5’-ATT GTG AAA ACC GAA CTT GGT CCA GAA GCA GTG AGC-3’ and 5’-TTC TGG ACC AAG TTC GGT TTT CAC AAT TTT GAA AGC-3’; and the peripheral primers 5’-GAG ACG GCC GAT GGT GAG CAA GGG CGA GGA GCT G-3’ and 5’-GAG AGG ATC CTT AAC TGT ACA ATA AAT AGA AG-3’. PCR fragments were cloned by ligation into the BamHI and EagI linearized pQCXIP vector.

The GFP-AH-MOSPD2 ΔCRAL-TRIO expression vector was obtained by PCR using GFP-MOSPD2 as template and the following central primers 5’-TGG ACC AAG CCA GGT TTT CAC AAT TTT GAA AGC AGC ATT CAT TAA CCA AGG CAT AGA ATT CGA AGC TTG AGC TCG AGA-3’ and 5’-TGG ACC AAG CCA GGT TTT CAC AAT TTT GAA AGC AGC ATT CAT TAA CCA AGG CAT AGA ATT CGA AGC TTG AGC TCG AGA-3’ and the peripheral primers 5’-GGA ATT GAT CCG CGG CCG CCG ATG GTG AGC AAG GGC GAG GAG CTGT-3’ and 5’-GGG CGG AAT TCC GGA TCT TAA CTG TAC AAT AAA TAG AAG AAA GAG GTG ACA AAA GCA AG-3’. PCR fragments were cloned using the SLiCE (Seamless Ligation Cloning Extract) method (Okegawa & Motohashi, 2015) into the NotI and BamHI linearized pQCXIP vector.

GFP-AH-VAP-A construct was obtained by PCR using the following primers: 5’-GGA ATT GAT CCG CGG CCG CCG ATG GTG AGC AAG GGC GAG GAG CTG T-3’, 5’-TGG ACC AAG CCA GGT TTT CAC AAT TTT GAA AGC AGC ATT CAT TAA CCA AGG CAT AGA ATT CGA AGC TTG AGC TCG AGA-3’, and 5’-TGC CTT GGT TAA TGA ATG CTG CTT TCA AAA TTG TGA AAA CCT GGC TTG GTC CAG CGT CCG CCT CAG GGG CCA TG-3’, 5’-GGG CGG AAT TCC GGA TCC TAC AAG ATG AAT TTC CCT AG-3’, and GFP-VAP-A as a template. PCR fragments were cloned by SLiCE into the NotI and BamHI linearized pQCXIP vector.

The GFP-MOSPD2-TM(SAC1) construct was obtained by PCR using the following primers: 5’-GGA ATT GAT CCG CGG CCG CCG ATG GTG AGC AAG GGC GAG GAG CTG T-3’, 5’-AAC AAC CAT GAT AAT AGG CAA AGC CAG GAA GAT ACA ACG CTG AAC TTG GTC TTC AAG CTT-3’, and 5’-TTC CTG GCT TTG CCT ATT ATC ATG GTT GTT-3’, 5’-GGG CGG AAT TCC GGA TCT CAG TCT ATC TTT TCT TTC TGG ACC AGT CT-3’, and GFP-MOSPD2 and GFP-TM(SAC1) as a template, respectively. PCR fragments were cloned by SLiCE into the NotI and BamHI linearized pQCXIP vector.

GFP-CRALTRIO-TM(SAC1) construct was obtained by PCR using the following primers: 5’-GGA ATT GAT CCG CGG CCG CCG ATG GTG AGC AAG GGC GAG GAG CTG T-3’, 5’-AAC AAC CAT GAT AAT AGG CAA AGC CAG GAA GGG GCC TTT AAA TAC ACT CAA TGG-3’, and 5’-TTC CTG GCT TTG CCT ATT ATC ATG GTT GTT-3’, 5’-GGG CGG AAT TCC GGA TCT CAG TCT ATC TTT TCT TTC TGG ACC AGT CT-3’, and GFP-MOSPD2 and GFP-TM(SAC1) as a template, respectively. PCR fragments were cloned using the SLiCE method into the NotI and BamHI linearized pQCXIP vector.

mScarlet-ER construct was obtained by PCR using the following primers: 5’-GGA ATT GAT CCG CGG CCG CCA CCA TGG TGA GCA AGG GC-3’, 5’-CCG GAC TTG TAC AGC TCG TCC ATG CCG CCG GTG GAG TGG CGG CCC TCG GAG C-3’, and 5’-GCT GTA CAA GTC CGG ATT CCT GGC TTT GCC TAT TAT CAT GGT TGT TGC CTT T-3’, 5’-GGG GGG GGC GGA ATT CTC AGT CTA TCT TTT CTT TCT GGA CCA GTC TGG GAG C-3’ and pmScarlet-C1 (gift from Dorus Gadella; Addgene plasmid # 85042; http://n2t.net/addgene:85042; RRID:Addgene_85042) and GFP-TM(SAC1) as a template, respectively. PCR fragments were cloned using the SLiCE method into the NotI and BamHI linearized pQCXIP vector.

mScarlet-MOSPD2 WT, mScarlet-MOSPD2 W201E and mScarlet-MOSPD2 RD/LD were obtained from GFP-MOSPD2 WT, GFP-MOSPD2 W201E and GFP-MOSPD2 RD/LD in which the GFP cassette was excised by SbfI and XhoI digestion and replaced using SLiCE by the coding sequence of mScarlet amplified by PCR using the primers: 5’-TGC ATT GGA ACG GAC CTG CAG CCA CCA TGG TGA GCA AGG GCG AGG CAG TGA TCA A-3’ and 5’-GCA GAA TTC GAA GCT TGA GCT CGA GAT CTG AGT CCG GAC TTG TAC AGC TCG TCC AT-3’ and pmScarlet-C1 as a template.

mScarlet-MOSPD2 ΔCRAL-TRIO was obtained from mScarlet-MOSPD2 WT in which the coding sequence of MOSPD2 was excised by XhoI and BamHI digestion and replaced using SLiCE by MOSPD2 ΔCRAL-TRIO coding sequence amplified by PCR using the primers: 5’-TCC GGA CTC AGA TCT CGA AGC TAT CCA CCA CTA GTA GAT GAT GAC TTC CAG ACC CCA CTG TGT GAG-3’ and 5’-GGG CGG AAT TCC GGA TCT TAA CTG TAC AAT AAA TAG AAG AAA GAG GTG ACA AAA GCA AG-3’.

All constructs were verified by DNA sequencing (Eurofins).

### Cell culture, transfection and infection

HeLa cells [American Type Culture Collection (ATCC) CCL-2, RRID:CVCL_0030] were maintained in DMEM (4.5 g/L glucose) with 5% fetal calf serum (FCS) and 40 μg/mL gentamycin. 293T cells (ATCC CRL-3216) were maintained in DMEM (4.5 g/L glucose) with 10% FCS, 100 UI/mL penicillin and 100 μg/mL streptomycin. Huh-7 cells (JCRB0403, RRID:CVCL_0336) were maintained in DMEM (4.5 g/L glucose) with 10% FCS, 0.1 mM NEAA, 1 mM sodium pyruvate and 40 μg/mL gentamycin. 501-MEL cells (RRID:CVCL_4633; obtained from Dr Colin Goding) were maintained in RPMI w/o HEPES with 10% FCS and 40 μg/mL gentamycin. MCF-7 cells (ATCC HTB-22) were maintained in DMEM (1 g/L glucose) with 10% FCS, 0.6 μg/mL insulin and 40 μg/mL gentamycin.

HeLa WT and Seipin-KO (kind gift from Hongyuan Robert Yang) were maintained in DMEM High Glucose (Dutscher) with 10 % Fetal Bovine Serum (FBS) and 1% Penicilin-Streptomycin. They were transfected using jetPEI transfection reagent (PolyPlus #101-10N).

Cells were transfected using X-tremeGENE 9 DNA Transfection Reagent (Roche). To generate retroviral particles, pQCXIP vectors were co-transfected with pCL-Ampho vector (Imgenex) into 293T retroviral packaging cell line. Retroviral infections were used to generate HeLa/Ctrl, HeLa/GFP-MOSPD2, HeLa/GFP-MOSPD2 RD/LD, HeLa/GFP-MOSPD2 ΔMSP, HeLa/GFP-MOSPD2 ΔTM, HeLa/GFP-MOSPD2 ΔCRAL-TRIO, and HeLa/mScarlet-ER cell lines. The HeLa/Ctrl cell line was obtained using the empty pQCXIP plasmid.

siRNA transfections were performed using Lipofectamine RNAiMAX (Invitrogen) according to the manufacturer’s instructions. Control siRNA (D-001810-10) and MOSPD2-targeting siRNAs (J-017039-09) were SMARTpool ON-TARGETplus obtained from Horizon Discovery.

Oleic acid was complexed with fatty acid-free BSA as described in Listenberger et al. (Listenberger & Brown, 2007), and diluted in cell culture medium. Unless otherwise stated, cells were treated overnight with 400 μM OA.

### CRISPR/Cas9-mediated genome editing

To generate MOSPD2 KO clones, HeLa cells were plated in 100 mm dishes and transfected with pX751 mCherry-Cas9 HF plasmid (gRNA deleting MOSPD2 exon 5: 5’-CAA GTG CAA CAG TTT CTC ATT-3’/5’-TGT TTG ACT ACA CTC ACA CT-3’) using X-tremeGENE 9 DNA Transfection Reagent (Roche). 48 hours after transfection, clones were sorted and isolated in 96-well plates using fluorescence-activated cell sorting (FACS, Fusion). Clones were then screened by PCR (5’-CAT CTT AGC TAC CAC CAC CTG AAC AGT TTA C-3’/ 5’-GCC TCG ACA TGC TAC CTC TCC-3’ and 5’-CAT CTT AGC TAC CAC CAC CTG AAC AGT TTA C-3’/5’-AAT TGC TGC TGA AGG GTT TGT AGG TAT C-3’), and further analyzed by Western Blot (anti-MOSPD2, HPA003334, Sigma-Aldrich, RRID:AB_2146004) and Sanger sequencing (Eurofins).

To generate endogenous mClover3-MOSPD2 knock-in cells, HeLa cells were plated in 100mm dishes and transfected with the pX852 plasmid (encoding mCherry-Cas9 and two gRNAs: 5’-AAC CGC AAT CAC ATC CAC GA-3’/5’-CAC CTC TGC CAT GAT CAC CG-3’) and the repair template (synthesized by ProteoGenix). The repair template was composed of two homology arms of 1000 bp flanking a puromycin resistance gene and mClover3 coding sequence separated by a P2A cleavage site. The insertion was made in the first exon of MOSPD2 genomic sequence to allow expression of a fusion protein with mClover3 at the N-terminus of MOSPD2. Three days after transfection, cells were selected with medium containing 0.5 μg/μL of puromycin. After 5 days of selection, cells were sorted in 96-well plates. Clones were screened by PCR (5’-GTG AAT TTT CAT GTA CAC TGG AGG ATG TTT GGC AGC-3’/5’-GCG AGG CGC ACC GTG GGC TTG TAC TCG GTC-3’ and 5’-ACA CAT GGC ATG GAC GAG CTG TAC AAG TCC-3’/ 5’-GCT TAA CTC CTT TCA CAG TAA CCA AAA TGA C-3’), and analyzed by Western Blot (anti-GFP and anti-MOSPD2) and Sanger sequencing (Eurofins).

### Immunofluorescence

Cells were grown on glass coverslips, fixed in 4% paraformaldehyde in PBS for 15 minutes, and permeabilized with 0.1% Triton X-100 in PBS for 10 minutes. After blocking with 1% bovine serum albumin in PBS (PBS-BSA), cells were incubated overnight at 4°C with the primary antibody in PBS-BSA. Primary antibodies were: rabbit anti-MOSPD2 (1:250; HPA003334, Sigma-Aldrich, RRID:AB_2146004), rabbit anti-GFP (1:1000; TP401, Torrey Pine Biolabs, RRID:AB_10013661), mouse anti-GFP (1:1000; 2A3, Euromedex) rabbit anti-Calnexin (1:1000; 10427-2-AP, Proteintech, RRID:AB_2069033), anti-Perilipin3 (1:1000; GP36, Progen), mouse anti-EEA1 (1:1000; 610457, BD Biosciences, RRID:AB_397830), rabbit anti-ORP1 (1:200; EPR8646, Abcam), rabbit anti-GM130 (1:500; 11308-1-AP, ProteinTech, RRID:AB_2115327), mouse anti-Lamp1 H4A3 (1:50; DSHB, RRID:AB_2296838) and mouse anti-OPA-1 (1:1000, 1A8, IGBMC). Cells were washed twice in PBS and incubated for 30 minutes with the secondary antibody (AlexaFluor 488 (RRID: AB_2535792 and AB_141607), AlexaFluor 555 (RRID: AB_2762848 and AB_162543), AlexaFluor 647 (RRID: AB_2536183 and AB_162542) from ThermoFisher Scientific and Abberior STAR 580 (RRID: AB_2620153) from Abberior. After two washes with PBS, the slides were mounted in ProLong Gold (Invitrogen). Observations were made with a Leica TCS SP5 inverted confocal microscope (63x, NA 1.4), a Leica SP8 UV inverted confocal microscope (63x, NA 1.4), and a spinning disk confocal microscope (CSU-X1, Nikon, 100x, NA 1.4). 2D-STED imaging was performed with a Leica SP8 STED 3X microscope in a thermostated chamber at 21°C and equipped with a STED motorized oil immersion objective (HC PL Apo 100X/ N.A. 1.40 CS2). Excitation was performed with white-light laser, depletion with a 775 nm pulsed laser (STED 775). Excitation and depletion lasers were calibrated with the STED auto beam alignment tool during imaging sessions. HeLa WT and KO Seipin cells were observed on a Carl ZEISS LSM 800 Airyscan microscope.

To stain LDs, cells were incubated after permeabilization with either BODIPY 493/503 (0.5 μg/ml in 150 mM NaCl; ThermoFisher), Nile Red (1:8000 in 150 mM NaCl; ThermoFisher) or HCS DeepRed LipidTOX (1:1000 in PBS; Invitrogen) for 20 minutes at room temperature.

### Quantification of ring-like/coma-shaped structures

HeLa WT cells were plated in 24-well plates (30,000 cells/well) and transfected the same day. Two days after transfection, cells were fixed using 4% PFA in PBS, washed two times with PBS and mounted on glass slides in ProLong Gold (Invitrogen).

Images were acquired on a Leica SP5 inverted confocal (63x oil objective, NA 1.4). The presence of enrichments (coma- and ring-shaped structures) was confirmed by eyes from two individuals.

### Quantification of LD number and size

Two million cells were plated in T75 flasks and allowed to grow for 48 hours. Thirty thousand cells were then plated in 24-well plates on glass coverslips. For the rescue experiment, HeLa MOSPD2 KO#1 cells were transfected the same day with different plasmids as previously described. After 48 hours, cells were fixed in 4% PFA PBS and permeabilized as described above. LDs were stained using BODIPY 493/503 and nuclei with Hoechst for 20 minutes. Cells were mounted on glass slides in ProLong Gold (Invitrogen). Images were acquired on a spinning disk CSU-X1 (Nikon, 100x NA 1.4), using the same setup every time (laser power, number of z-slices, exposition length). Cells were selected based on the nuclei channel to avoid any bias.

For image processing (illustrated in Fig. S4A), a z-stack projection (max intensity) was performed on ImageJ Fiji (Schindelin *et al*, 2012). These images were then processed using CellProfiler (McQuin *et al*, 2018). First, cells were manually segmented to create cell masks. Then, LDs were identified as objects with a diameter ≥ 2 pixels (*i.e*. 220 nm) within cell masks using the global threshold strategy and the minimum cross-entropy method. Multiple parameters (object intensity, object neighbors and object size/shape) were analyzed on identified LDs and these data were treated using Spyder 4.1 (Python 3.7) and GraphPad Prism.

### Quantification of ER fluorescence signal around LDs

Stable mScarlet-ER HeLa cells were plated in 24-well plates (30,000 cells/well) and transfected the same day with either GFP-VAP-A or GFP-MOSPD2. Thirty-six hours later, cells were treated with 50 μM of OA for 6 hours. Cells were fixed in PFA 4% PBS, permeabilized with Triton X-100 0.1% and LDs were stained using HCS DeepRed LipidTOX as described above. Images were acquired on a Leica SP5 inverted confocal (63x oil objective, NA 1.4) and cells were selected based on the GFP signal.

Image processing (illustrated in Fig. S4B) was performed on CellProfiler. In brief, cells were manually selected and the nucleus was excluded (cell mask). The LDs were identified as objects superior or equal to 4 pixels of diameter using the global threshold strategy and the minimum cross-entropy method. Then, 2 pixels-wide area were added from 0 to 20 pixels around LDs. These areas are mutually exclusive, meaning that the same pixel can be measured only once (*i.e*. for one LD). Multi-parametric measurements were performed for each area around LDs in the GFP channel (MOSPD2 and VAP-A) and mScarlet channel (ER marker). Data were analyzed on Excel and GraphPad.

### Correlative Light Electron Microscopy (CLEM)

Electron microscopy was performed as previously described (Di Mattia *et al*, 2018; Wilhelm *et al*, 2017; Alpy *et al*, 2013). Cells grown on carbon-coated sapphire disks were cryoprotected with DMEM containing 10% FCS and frozen at high pressure (HPM 10 Abra Fluid AG). Samples were then freeze-substituted and embedded in lowicryl HM20. Thick sections (^~^250 nm) were collected on carbon-coated copper grids (200 Mesh; AGS160; Agar Scientific).

EM grids were placed on a MatTek glass bottom dish in a drop of water and imaged with a spinning disk confocal microscope (CSU-X1, Nikon, 100x oil objective, NA 1.4). The position of the imaged cells was determined using the asymmetrical center mark of the grid. Then, samples were imaged with a transmission electron microscope (Philips CM12) coupled to an Orius 1000 CCD camera (Gatan). Images were processed and merged with the open-source software Icy (de Chaumont *et al*, 2012) using the eC-CLEM plugin (Paul-Gilloteaux *et al*, 2017).

### Fluorescence Recovery After Photobleaching (FRAP)

Cells were plated on 35 mm glass bottom dishes (MatTek), transfected with plasmids encoding GFP-MOSPD2 or GFP-Plin2, and allowed to grow for 24 hours. Cells were then treated with 400 μM oleic acid overnight. HCS LipidTOX Deep Red Neutral Lipid Stain (H34477, Invitrogen) at a 1:2000 dilution was added to the medium without phenol red 10 minutes prior imaging. Experiments were performed using the spinning disk CSU-X1 (Nikon, 100x oil objective, NA 1.4). A region of interest was photobleached with the 405 nm laser line at 15% laser power and 5 repetitions. Recovery of fluorescence was monitored every second for 1 minute immediately after photobleaching.

### Protein expression and purification

The recombinant WT and mutant CRAL-TRIO_His6_ [MOSPD2 2-246] and MSP_His6_ [MOSPD2 315-445] proteins, and the recombinant MOSPD2_His6_ [MOSPD2 1-490] protein corresponding to the full-length protein with its carboxyl-terminal transmembrane region deleted, were expressed in *E. coli* BL21 (DE3) at 20°C for 16 hours upon induction with 1 mM IPTG (at an optical density OD_600nm_ = 0.5). Cells were suspended in lysis buffer [50 mM sodium phosphate pH 8.0, 300 mM NaCl, 10 mM Imidazole, protease inhibitors tablets (cOmplete, Roche)]. Cells were lysed by a Cell Disruptor TS SERIES (Constant Systems Ltd), and the lysate was first centrifuged at 3,500 g for 15 minutes, then at 50,000 x g for 45 minutes, and filtered through a 0.22 μm membrane. Purification was performed on an ÄKTA Start chromatography system (GE Healthcare Life Sciences) using HisTrap HP 1 mL columns. Proteins were eluted with Elution buffer [20 mM sodium phosphate pH 7.4, 250 mM Imidazole], and further purified by gel filtration (HiLoad 16/60 Superdex 200; GE) in GF Buffer (20 mM Tris-HCl pH 7.5, 150 mM NaCl). Proteins were concentrated with an Amicon Ultra-15 10 kDa centrifugal filter unit (Merck). Protein concentration was determined by UV-spectroscopy.

### Peptide synthesis

Peptides were synthesized on an Applied biosystem 433A peptide synthesizer using standard Fmoc chemistry, and purified by reverse phase HPLC using a preparative scale column (Phenomenex: Kinetex EVO C18, 100 Å, 5 μM, 250 × 21.2 mm). Molecular weight and purity of the peptides were confirmed by mass spectrometry.

### SDS-PAGE, Western blot and Coomassie Blue staining

SDS-PAGE and Western blot analysis were performed as previously described (Alpy *et al*, 2005) using the following antibodies: rabbit anti-GFP (1:2000; TP401, Torrey Pine Biolabs), rabbit anti-MOSPD2 (1:250; HPA003334, Sigma-Aldrich), rabbit anti-mCherry (1:1000; ab167453, Abcam) and mouse anti-actin (1:5000; ACT-2D7, Euromedex).

Protein gels were stained with Coomassie blue (PageBlue Protein Staining Solution; Thermo Fisher Scientific).

### Lipids

DOPC (1,2-dioleoyl-*sn*-glycero-3-phosphocholine), DOPE (1,2-dioleoyl-sn-glycero-3-phosphoethanolamine), diphytanoyl-PC (1,2-diphytanoyl-*sn*-glycero-3-phosphocholine), DOPS (1,2-dioleoyl-*sn*-glycero-3-phospho-L-serine), diphytanoyl-PS (1,2-diphytanoyl-*sn*-glycero-3-phospho-L-serine), NBD-PE (1,2-dioleoyl-*sn*-glycero-3-phosphoethanolamine-N-(7-nitro-2-1,3-benzoxadiazol-4-yl)), 18:1 Liss Rhod PE (1,2-dioleoyl-*sn*-glycero-3-phosphoethanolamine-N-(lissamine rhodamine B sulfonyl)) and DOGS-NTA-Ni^2+^ (1,2-dioleoyl-*sn*-glycero-3-[(N-(5-amino-1-carboxypentyl) iminodiacetic acid) succinyl]) were purchased from Avanti Polar Lipids. Glyceryl trioleate was purchased from Sigma-Aldrich

### Thin Layer Chromatography (TLC)

Two million cells were plated in T75 flask and allowed to grow for 48 hours. Then, 500,000 cells were plated in 6-well plates. After 24 hours, cells were lysed with 0.1% SDS. Proteins were quantified and the lysate equivalent to 450 μg of proteins was transferred in a new tube and the volume adjusted to 800 μL. Lipids were extracted using Bligh & Dyer protocol (water/chloroform/methanol with a ratio of 1.8:2:2). After 5 minutes of centrifugation at 400 g, the lower-phase containing the lipids was transferred into a disposable glass tube and dried under a nitrogen stream. Lipids were solubilised in three drops of a chloroform/methanol mix (9:1) and applied to a HPTLC plates by capillarity. Cholesteryl ester (CE) and triacylglycerol (TAG) standards were also applied to the plates. Plates were then developed in a neutral lipid solvent (hexane/diethylether/AcOH with a ratio of 80:20:2) and primuline solution was used for revelation. Images were acquired on an ImageQuant LAS 4000 mini (GE Healthcare Life Sciences) using EtBr as fluorescence and the 605DF40 filter. The intensity of CE and TAG bands were quantified using Fiji.

### Enzymatic quantification of lipids

Two million cells were plated in T75 flask and allowed to grow for 48 hours. Then, 500 000 cells were plated in 6-well plates. After 24 hours, cells from 2 wells were scraped in PBS and lipids extracted using the same protocol as for TLC (Bligh & Dyer protocol (Bligh & Dyer, 1959)). Lipids were solubilised in 200 μL of ethanol 95%. Quantifications were then performed according to the manufacturers’ instructions (Cholesterol Amplex Red from Thermofisher (A12216), High Sensitivity Triglyceride Fluorometric Assay Kit from Sigma (MAK264-1KT) and Phospholipid quantification Assay Kit from Sigma (CS0001-1KT)).

### Artificial lipid droplets (aLDs) preparation

Artificial LDs were prepared as described in Prévost *et al*. (2018) with slight modifications. Phospholipids (DOPC, DOPE and Rhodamine-DOPE with a ratio of 73:25:2 respectively) were added to triolein at a 0.5% molar ratio in a glass tube. The solvent was evaporated under a stream of nitrogen and 1 mL of HK buffer (50 mM Hepes pH 7.2, 120 mM potassium acetate) was added. The solution was vortexed for 3 minutes. Then, aLDs were extruded 11 times through polycarbonate filters (Nuclepore Track-Etch Membrane, Whatman) with a pore diameter of 100 nm using a mini-extruder (Avanti Polar Lipids). The size was verified by dynamic light scattering measurements on a DynaPro (Protein Solutions). The preparation was used in a couple of hours after extrusion.

### Artificial LDs (aLDs) flotation assay

Each peptide (1 μM) was mixed with aLDs (1 mM), vortexed and incubated at room temperature for 15 minutes. The solution (volume of 390 μL) was adjusted to 30% (w/w) sucrose by mixing with 260 μL of a 75% sucrose HK buffer. Two layers (520 μL of 25% sucrose and 130 μL of sucrose-free HK buffer) were gently added on top. Samples were centrifuged at 240,000 x g for 1 hour in a swing rotor (SWTi 60) at 20°C and decelerated without brake. The bottom (520 μL), middle (520 μL) and top (260 μL) fractions were collected and the fluorescence was measured using a fluorimeter (Pherastar FSX, BMG LABTECH).

### aLDs and peptide interaction assay

Artificial LDs (aLDs) were prepared following the protocol developed in Chorlay et al (Chorlay & Thiam, 2020). Briefly, 5 μL of triolein were mixed with 70μL of HKM buffer (50 mM Hepes, 120 mM K-acetate, and 1 mM MgCl2 at pH 7.4) and vortexed for 5 seconds and sonicated in a bath sonicator for 10 seconds. Then each peptide was separately mixed and incubated with aLDs at a concentration of 1μM. The resulting emulsions were then introduced in a glass chamber for visualization. Fluorescence data were acquired 30 minutes after incubation, with a laser scanning microscope (LSM 800, Carl Zeiss). Fluorescence intensity was measured after segmentation of individual aLDs.

### aLDs pull-down assay

aLDs pull-down was performed as described in Kassas et al. (Kassas *et al*, 2017). NTA-Ni^2+^ beads (PureProteome Nickel Magnetic Beads, LSKMAGH10, Millipore) were first washed with HK buffer. Then, 1 μM of recombinant proteins (MSP_His6_, WT and W201E CRAL-TRIO_His6_) was added to the beads and incubated 20 minutes under agitation at 4°C. To remove the excess of proteins, beads were washed twice with HK buffer. Afterwards, aLDs (1 mM) were added to the beads and incubated 20 minutes again, under agitation at 4°C. Beads were then washed 3 times with HK buffer and resuspended in a final volume of 30 μL of HK buffer. For imaging, 10 μL of the suspension were dropped on a glass bottom dish (MatTek) and imaged on a spinning disk CSU-X1 (Nikon, 100x NA 1.4). Fluorescence was measured with a fluorimeter (Pherastar FSX, BMG LABTECH) using 20 μL of suspension.

### Liposomes preparation

Lipids stored in stock solutions in CHCl3 were mixed at the desired molar ratio in glass tubes. The solvent was removed in a dryer-block at 33°C under a nitrogen flow. If the mixture contained DOGS-NTA-Ni^2+^, it was pre-warmed at 33°C for 5 minutes prior to drying. The lipid film was hydrated in 50 mM Hepes, pH 7.2, 120 mM K-Acetate (HK) buffer to obtain a suspension of multi-lamellar vesicles. The multi-lamellar vesicles suspensions were frozen and thawed five times and then extruded through polycarbonate filters of 0.2 μm pore size using a mini-extruder (Avanti Polar Lipids). Liposomes were stored at 4°C and in the dark when containing fluorescent lipids and used within 2 days.

### Flotation experiment

The CRAL-TRIO_His6_ protein (0.75 μM) was incubated with liposomes (0.75 mM total lipid) with a given lipid composition doped with 0.1 mol% NBD-PE in 150 μL of HK buffer at 25°C for 10 minutes under agitation at 800 rpm. The suspension was adjusted to 28% (w/w) sucrose by mixing 100 μL of a 60% (w/w) sucrose solution in HK buffer and overlaid with 200 μL of HK buffer containing 24% (w/w) sucrose and 50 μL of sucrose-free HK buffer. The sample was centrifuged at 240,000 × g in a swing rotor (TLS 55 Beckmann) for 1 hour. The bottom (250 μL), middle (150 μL) and top (100 μL) fractions were collected. The bottom and top fractions were analyzed by SDS-PAGE by direct fluorescence and after staining with SYPRO Orange, using a FUSION FX fluorescence imaging system.

### Circular dichroism

The experiments were performed on a Jasco J-815 spectrometer at room temperature with a quartz cell of 0.05 cm path length (Starna). The CRAL-TRIO_His6_ protein (WT or W201E mutant) was dialysed against a 20 mM Tris, pH 7.4, 120 mM NaF buffer to remove glycerol. Each spectrum is the average of ten scans recorded from 185 to 260 nm with a bandwidth of 1 nm, a step size of 0.5 nm and a scan speed of 50 nm.min^−1^. The protein concentration was 6.7 μM. A control spectrum of buffer was subtracted from each protein spectrum. The spectra were analyzed in the 185-240 nm range using the BeStSel method provided on-line (Micsonai *et al*, 2015).

### Dynamic light scattering measurements of liposome aggregation

The experiments were performed at 25°C in a Dynamo apparatus (Protein Solutions). L_A_ liposomes (50 μM total lipids) made only of DOPC or composed of DOPC/DOGS-NTA-Ni^2+^ (98/2 mol/mol) were mixed with L_B_ liposomes (50 μM, made of DOPC or composed of diphytanoyl-PC and diphytanoyl-PS (70/30 mol/mol)) in 20 μL of a freshly degassed HK buffer in a quartz cell. A first set of 12 autocorrelations curves was acquired to determine the size distribution of the initial liposome suspensions. Then, MOSPD2_His6_ or MSP_His6_ (0.4 μM final concentration) was added manually and mixed thoroughly. The kinetics of aggregation was measured during 23 minutes by acquiring one autocorrelation curve every 10 seconds. At the end of the experiment, a set of 12 autocorrelation functions was acquired. The data were analyzed using two different algorithms provided by the Dynamics v6.1 software (Protein Solutions). During the kinetics, the autocorrelation functions were fitted assuming that the size distribution is a simple Gaussian function. This mode, referred to as the monomodal or cumulant algorithm, gives a mean hydrodynamic radius, RH, and the width (or polydispersity). Before and after the aggregation process, the autocorrelation functions were fit-ted using a more refined algorithm, referred as a regularization algorithm. This algorithm is able to resolve several populations of different sizes, such as free liposomes and liposome aggregates.

### Statistical analyses

Statistical analyses were performed using the Mann-Whitney, or the Kruskal-Wallis non-parametric test, and with the One-way ANOVA parametric test (Prism, GraphPad). Conditions were compared with the Dunn’s and Tukey’s multiple comparisons tests, respectively. P-values <0.05, <0.01, <0.001 and <0.0001 are identified with 1, 2, 3 and 4 asterisks, respectively, ns: p≥0.05.

## Acknowledgments

We thank the members of the Molecular and Cellular Biology of Breast Cancer team (IGBMC) for helpful advice and discussions. We are grateful to the members of the IGBMC Imaging Center (ICI), especially Elvire Guiot for her help with FRAP analysis, and Bertrand Vernay for his help with CellProfiler. We thank Paolo Ronchi from the Electron Microscopy Core Facility at the EMBL Heidelberg, and Coralie Speigelhalter and Danièle Spehner from the IGBMC for their help with electron microscopy. We thank Marko Lampe from the Advanced Light Microscopy Facility at the EMBL Heidelberg for his help with STED microscopy. We thank the IGBMC cell culture facility (Betty Heller), the peptide synthesis facility (Pascal Eberling), the Flow Cytometry facility (Claudine Ebel and Muriel Philipps), Molecular Biology and Virus Service (Paola Rossolillo and Nicole Jung), the Integrated Structural Biology platform (Catherine Birck), and the polyclonal and monoclonal antibody facility (Mustapha Oulad-Abdelghani) for their excellent technical assistance.

M.Z. received a fellowship from ITMO Cancer AVIESAN (Alliance Nationale pour les Sciences de la Vie et de la Santé / National Alliance for Life Sciences and Health) within the framework of the Cancer Plan (https://itcancer.aviesan.fr/). T.D.M. received a fellowship from the Fondation pour la Recherche Médicale (https://www.frm.org/). A.M. received a fellowship from the Fondation ARC pour la recherche sur le cancer (https://www.fondation-arc.org/). J.E. received a fellowship from IMCBio.

This work was supported by grants from the Agence Nationale de la Recherche ANR (grant ANR-21-CE13-0014-01; https://anr.fr/), and from the Ligue Contre le Cancer (Conférence de Coordination Interrégionale du Grand Est; https://www.ligue-cancer.net), SEVE Sein et Vie. We also acknowledge funds from the Institut National de Santé et de Recherche Médicale (http://www.inserm.fr/), the Centre National de la Recherche Scientifique (http://www.cnrs.fr/), the Université de Strasbourg (http://www.unistra.fr) and the grant ANR-10-LABX-0030-INRT, a French State fund managed by the Agence Nationale de la Recherche under the frame program Investissements d’Avenir ANR-10-IDEX-0002-02.

