## Supplementary figures for "MOSPD2 a new endoplasmic reticulum-lipid droplet tether functioning in LD homeostasis"

### Figure S1

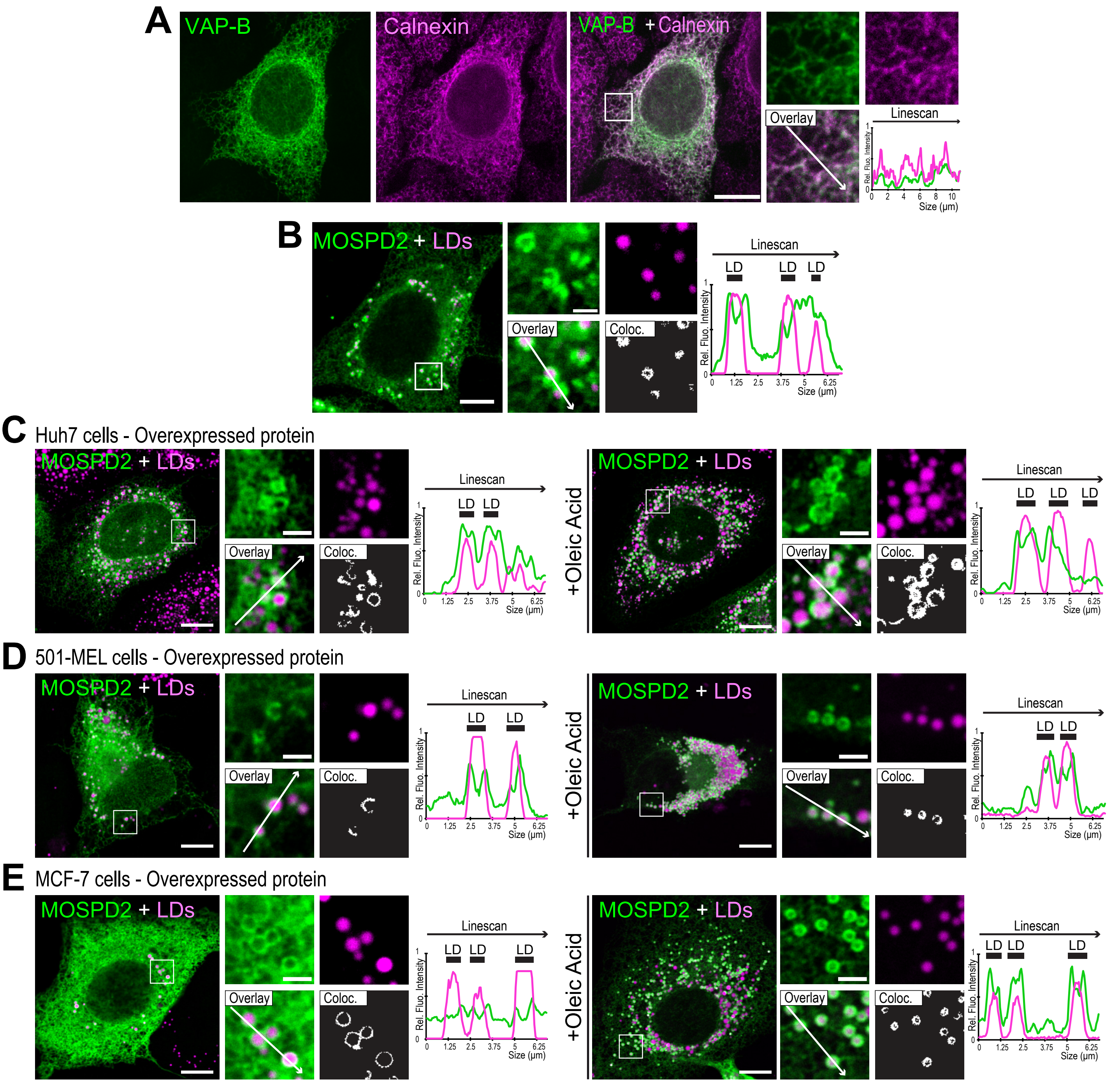


**Figure S1: MOSPD2 is enriched around LDs in different cell lines.**
A. HeLa cells expressing GFP-VAP-B (green) were labeled with anti-Calnexin antibodies (magenta). Spinning disk confocal microscope (Nikon CSU-X1, 100x NA 1.4) images.
B. HeLa cells expressing GFP-MOSPD2 (green) and not treated with OA were labeled with Nile Red to stain LDs (magenta). Confocal microscope (Leica SP5, 63x NA 1.4) images.
C, D, E. Colocalization in Huh-7 (C), 501-MEL (D) and MCF7 (E) of GFP-MOSPD2 (green). Cells were either treated with OA (right) or not treated (left). LDs were stained using Nile Red (magenta).

Data information C, D, and E: Subpanels on the right are higher magnification images of the outlined areas. The overlay panel shows merged channels. The coloc panel displays a colocalization mask in which pixels of the green and magenta channels that co-localize are shown in white. Linescan shows fluorescence intensities of the green and magenta channels along the white arrow from the overlay subpanel. Black rectangles indicate the position of Lipid Droplets (LD). Scale bars: 10 µm (insets 2 µm). Confocal microscope (Leica SP5, 63x NA 1.4) images.

### Figure S2

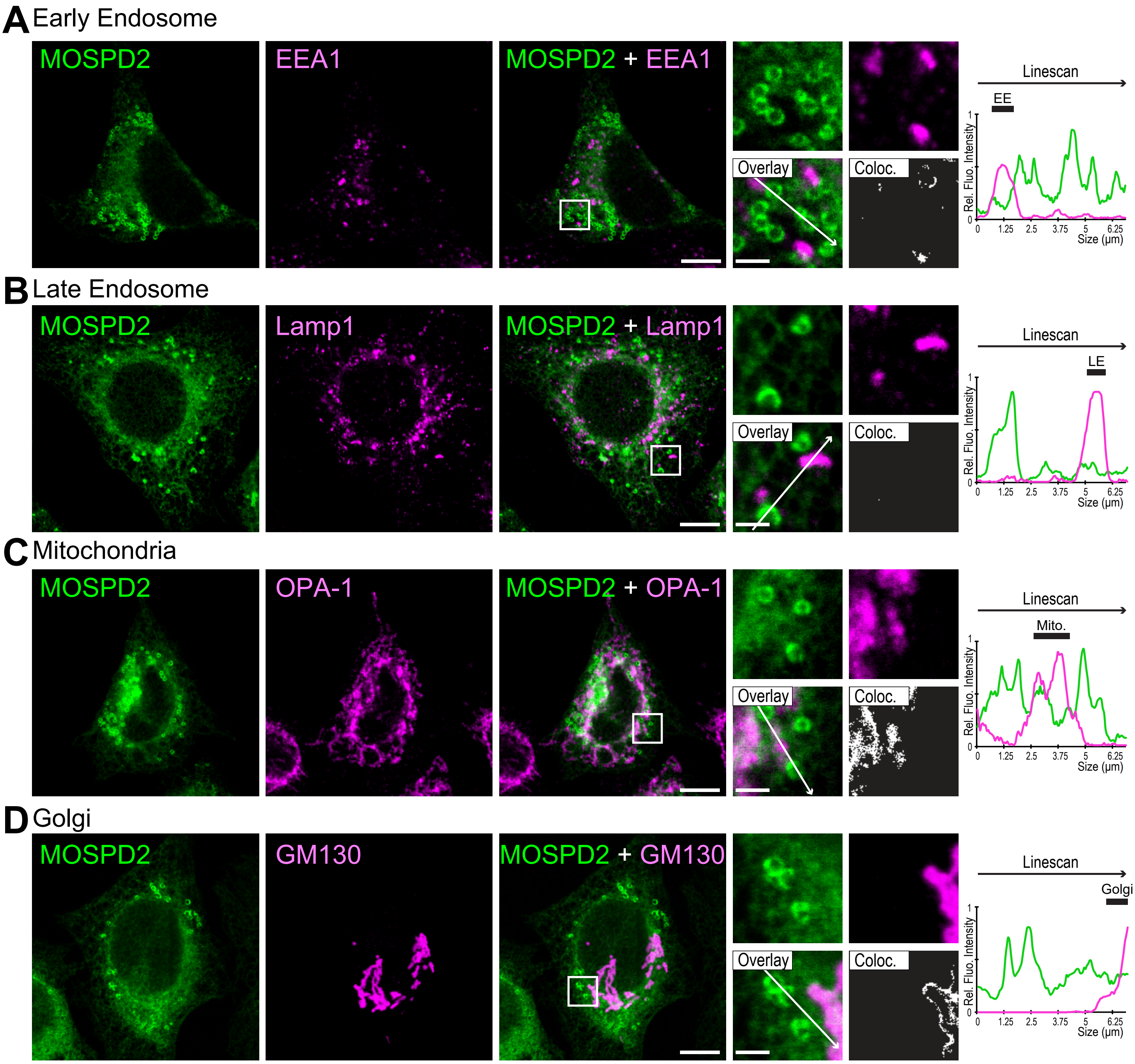


**Figure S2: Colocalization of MOSPD2 with different organelles.**
A-D. Colocalization in HeLa cells transfected with GFP-MOSPD2 (green) of MOSPD2 and EEA1 (A, magenta), Lamp1 (B, magenta), OPA-1 (C, magenta) and GM130 (D, magenta).
Data information: Subpanels on the right are higher magnification images of the outlined areas. The overlay panel shows merged channels. The coloc panel displays a colocalization mask in which pixels of the green and magenta channels that co-localize are shown in white. Linescan shows fluorescence intensities of the green and magenta channels along the white arrow from the overlay subpanel. Black rectangles indicate the position of Early Endosomes (EE; in A), Late Endosome (LE; in B), Mitochondria (Mito.; in C) and the Golgi apparatus (Golgi; in D). Scale bars: 10 µm (insets 2 µm). Images were acquired on a confocal microscope (Leica SP5; 63x NA 1.4).

### Figure S3

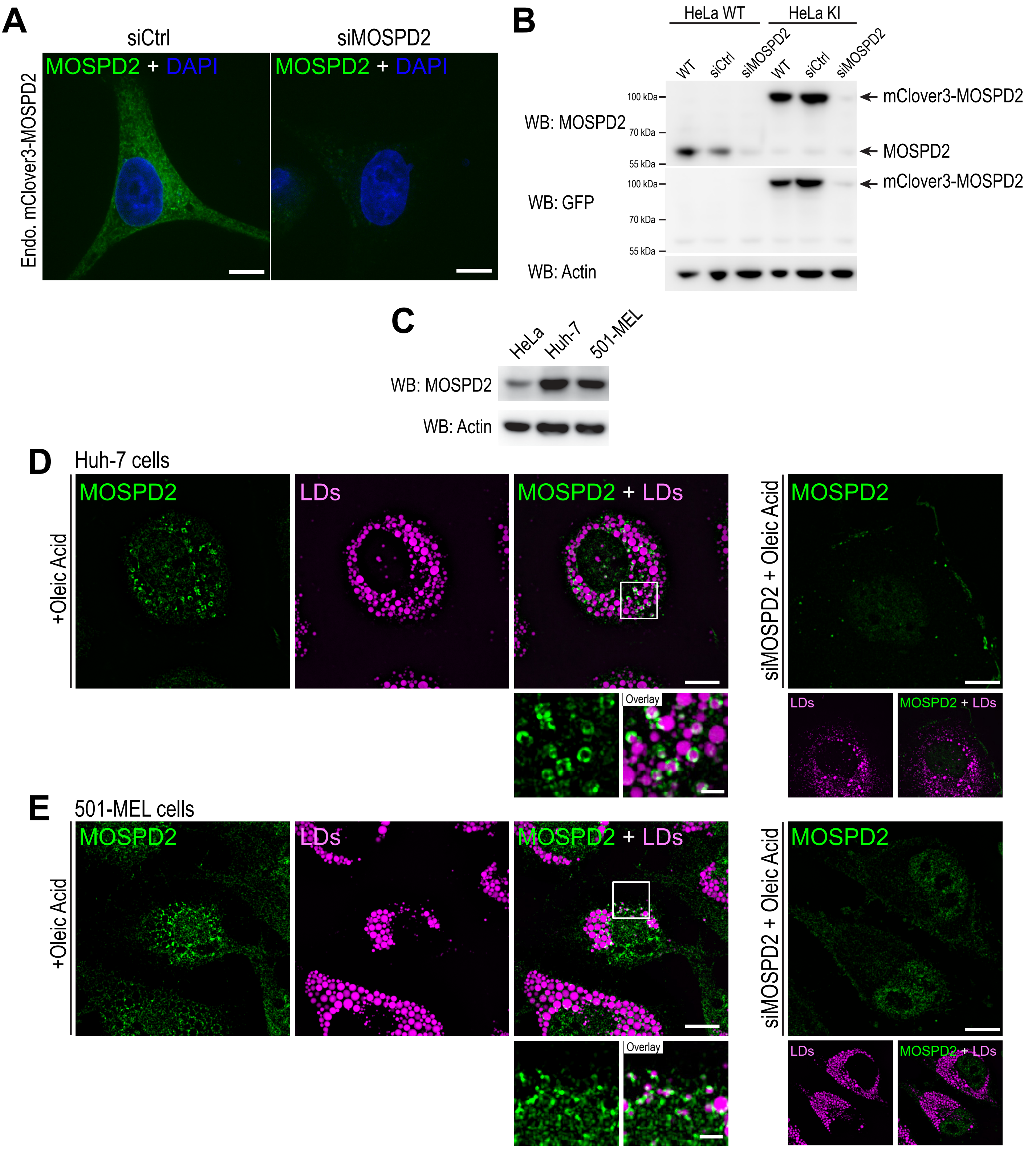


**Figure S3: Characterization of CRISPR/Cas9 knock-in HeLa cells and endogenous localization of MOSPD2.**

A. Confocal images of live HeLa cells expressing mClover3-MOSPD2 (green) at the endogenous level and transfected with control siRNAs (siCtrl) and siRNAs targeting MOSPD2 (siMOSPD2) to confirm the specificity of mClover3 signal. Scale bars: 10 µm. Images were acquired on a spinning disk confocal microscope (Nikon CSU-X1, 100x NA 1.4).

B. Western Blot analysis of MOSPD2 expression in WT (HeLa WT) and mClover3-MOSPD2 knock-in (HeLa KI) HeLa cells transfected with control siRNAs (siCtrl) and siRNAs targeting MOSPD2 (siMOSPD2). mClover3 was detected using anti-GFP antibodies.

C. Western Blot analysis of MOSPD2 expression in HeLa, Huh-7 and 501-MEL cells.

D and E. Colocalization of endogenous MOSPD2 (labeled with anti-MOSPD2, in green) and LDs (labeled with LipidTOX, in magenta) in Huh-7 cells (D) or 501-MEL (E) after OA treatment. Panels on the right show the aspecific signal of anti-MOSPD2 antibodies in cells transfected with siRNAs targeting MOSPD2 (siMOSPD2). Images were acquired on a spinning disk confocal microscope (Nikon CSU-X1, 100x NA 1.4). Subpanels on the right are higher magnification images of the area outlined. The overlay panel shows merged channels. Scale bars: 10 µm (insets 2 µm).

### Figure S4

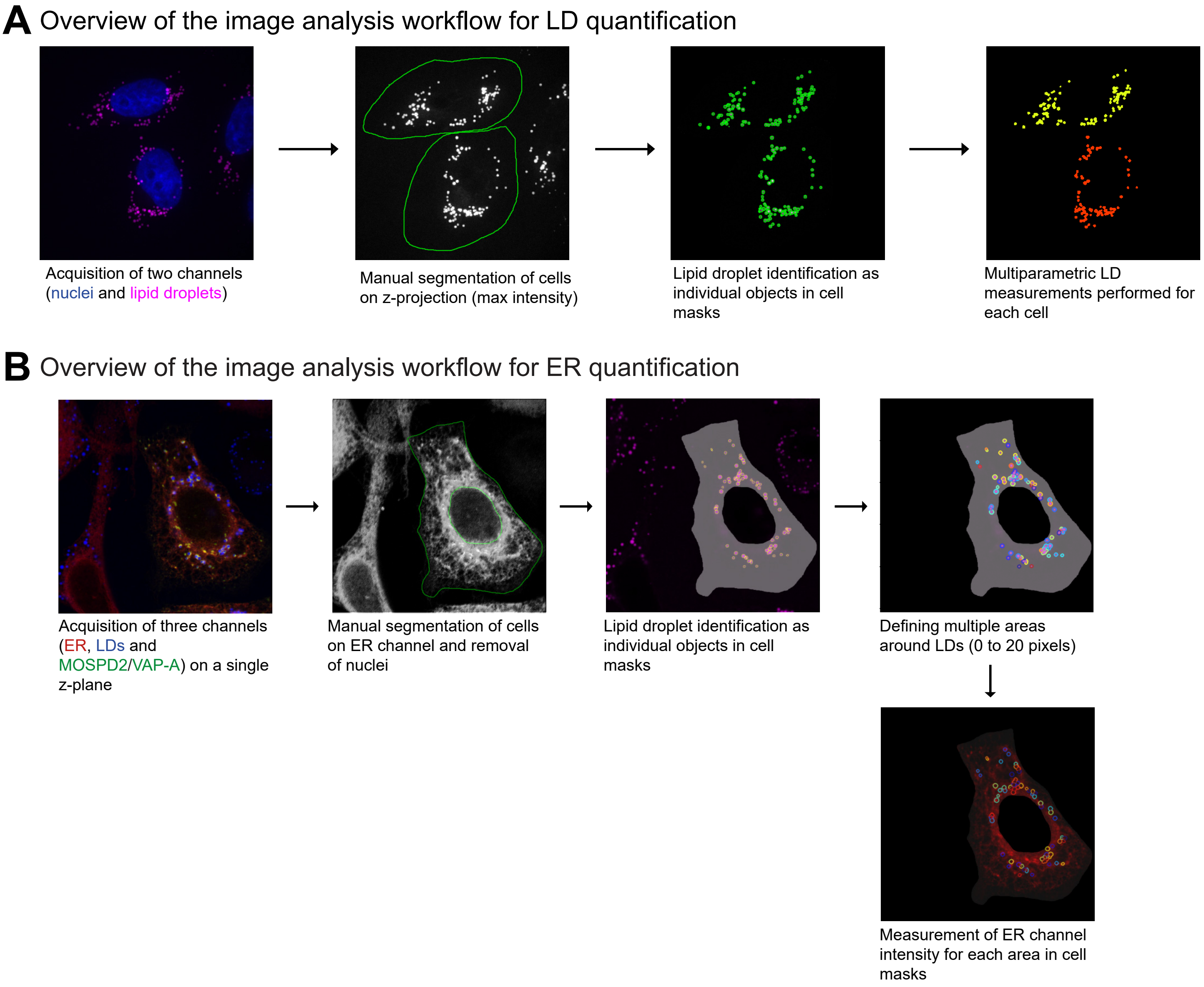


**Figure S4: Image analysis workflows for LD and ER quantifications.**
A. Cells stained with BODIPY 493/503 (LDs) and Hoechst (nuclei) were imaged on multiple z slices using a confocal microscope. A z-stack projection (max intensity) image was generated using Fiji and processed using CellProfiler. Cells were manually segmented and LDs identified as objects ≥ 2 pixels of diameter. Multi-parametric object measurements were performed on the identified LDs.
B. Cells were treated with oleic acid at 50 µM for 6 hours before imaging. Three channels were acquired: nuclei (stained with Hoescht), ER marker (mScarlet-ER) and MOSPD2/VAP-A (tagged with GFP). Cells were manually segmented and masks of the cytoplasm (*i.e.* without the nuclei) were generated with CellProfiler. LDs were identified as objects ≥ 4 pixels of diameter. Multiple areas (2-pixel wide) from 0 to 20 pixels around each LD were added. Multi-parametric measurements were performed for each area around LDs in the red (mScarlet-ER) and green (GFP-MOSPD2/VAP) channels.

### Figure S5

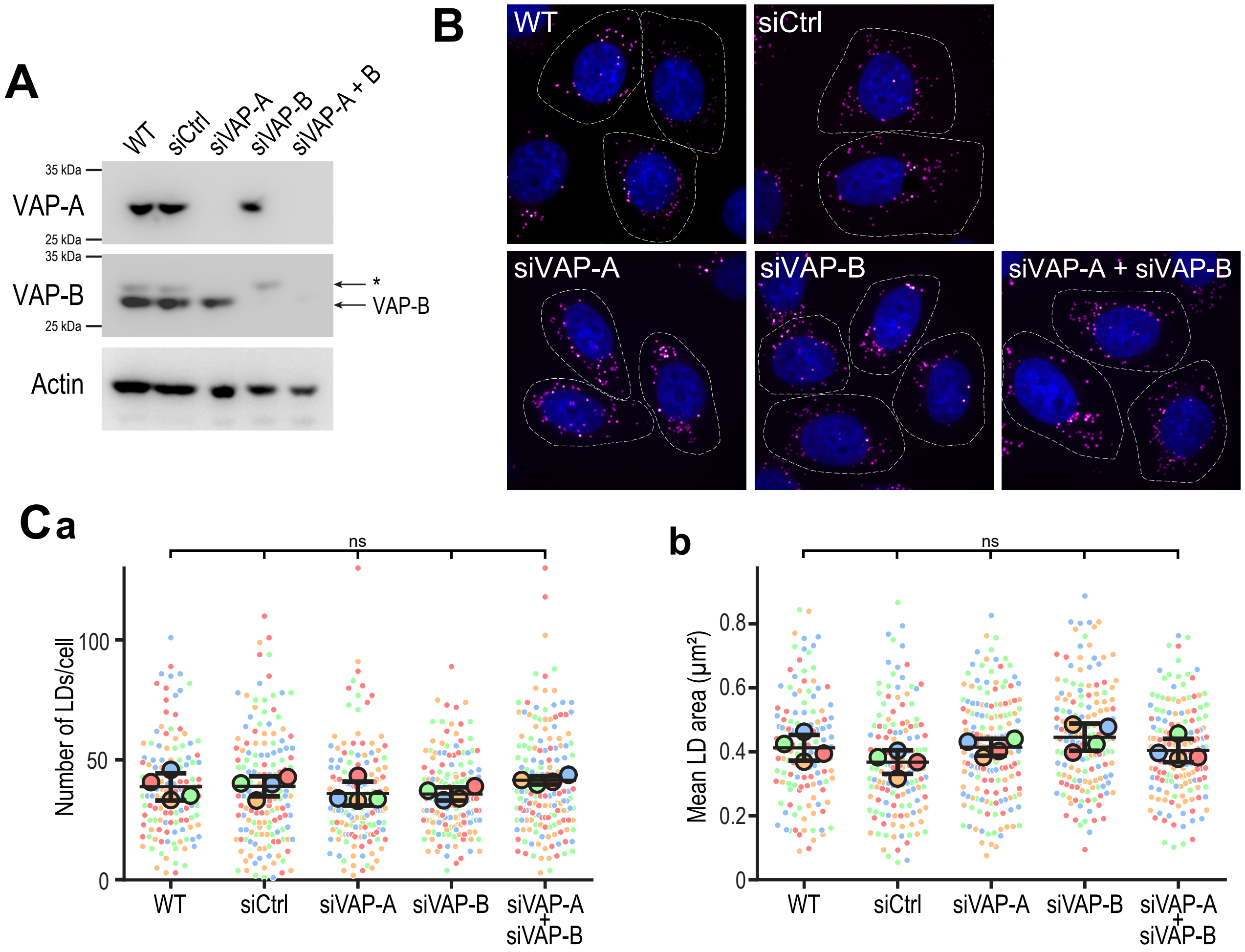


**Figure S5: VAP-A and VAP-B are not involved in LD homeostasis.**

A. Western blot analysis of VAP proteins level in control HeLa cells (WT), HeLa cells transfected with control siRNAs (siCtrl), and with siRNAs targeting VAP-A (siVAP-A), VAP-B (siVAP-B) or both (siVAP-A + B). Bands annoted with a * on VAP-B blot correspond to VAP-A signal.

B. Representative confocal images of parental HeLa cells (WT) and HeLa cells transfected with control siRNAs (siCtrl), and with siRNAs targeting VAP-A (siVAP-A), VAP-B (siVAP-B) or both (siVAP-A + B). Cells were labeled with BODIPY 493/503 (LD, magenta) and Hoechst 33258 (nuclei, blue). Scale bars: 10 µm. Images were acquired on a spinning disk confocal microscope (Nikon CSU-X1, 100x NA 1.4).

C. Number (left) and area (right) of LDs in cells shown in B. Data are displayed as Superplots showing the mean number (left) and area (right) of LDs per cell (small dots), and the mean number (left) and area (right) of LDs per independent experiment (large dots). Independent experiments (n = 4) are color-coded. Means and error bars (SD) are shown as black bars. Data were collected from 98 (WT), 118 (siCtrl), 134 (siVAP-A), 129 (siVAP-B) and 135 (siVAP-A + VAP-B) cells. One-way ANOVA with Tukey’s multiple comparisons test (ns: not significant; n = 4 independent experiments).

### Figure S6

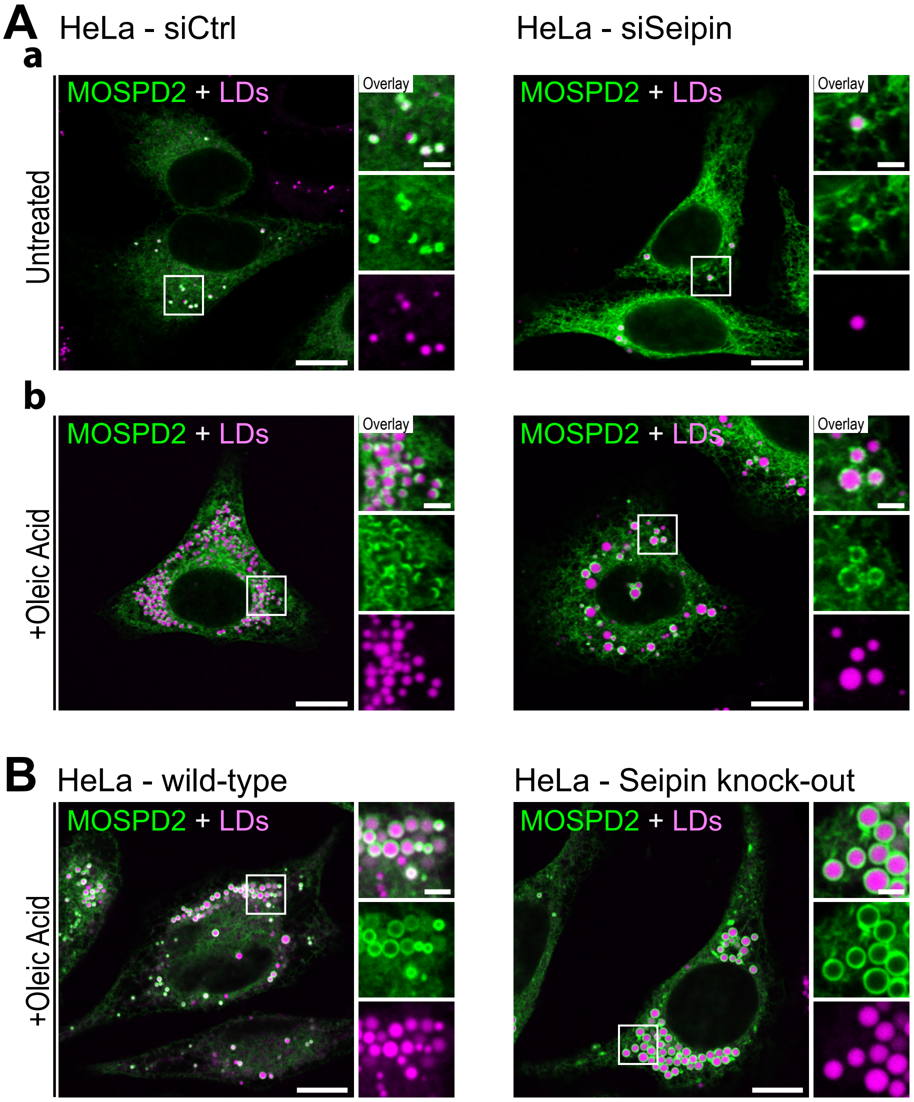


**Figure S6: Seipin is dispensable for MOSPD2-mediated ER-LD contact formation.**

A: Representative confocal images of the GFP-MOSPD2 WT (green) localization in cells transfected with control siRNAs (left) and siRNAs targeting Seipin (right) and left untreated (a) or treated with OA (b). LDs were stained with Nile Red (magenta). Confocal microscope (Leica SP8; 63x NA 1.4) images.

B:Representative confocal images of the GFP-MOSPD2 WT (green) localization in wild-type (left) and Seipin knock-out (right) cells treated with OA. LDs were stained with LipidTox (magenta).

Scale bars: 10 µm (insets 2 µm). Note that Seipin silencing or knock-out results in heterogeneous lipid droplet size. In absence of Seipin, MOSPD2 still mediates ER-LD contact formation.

### Figure S7

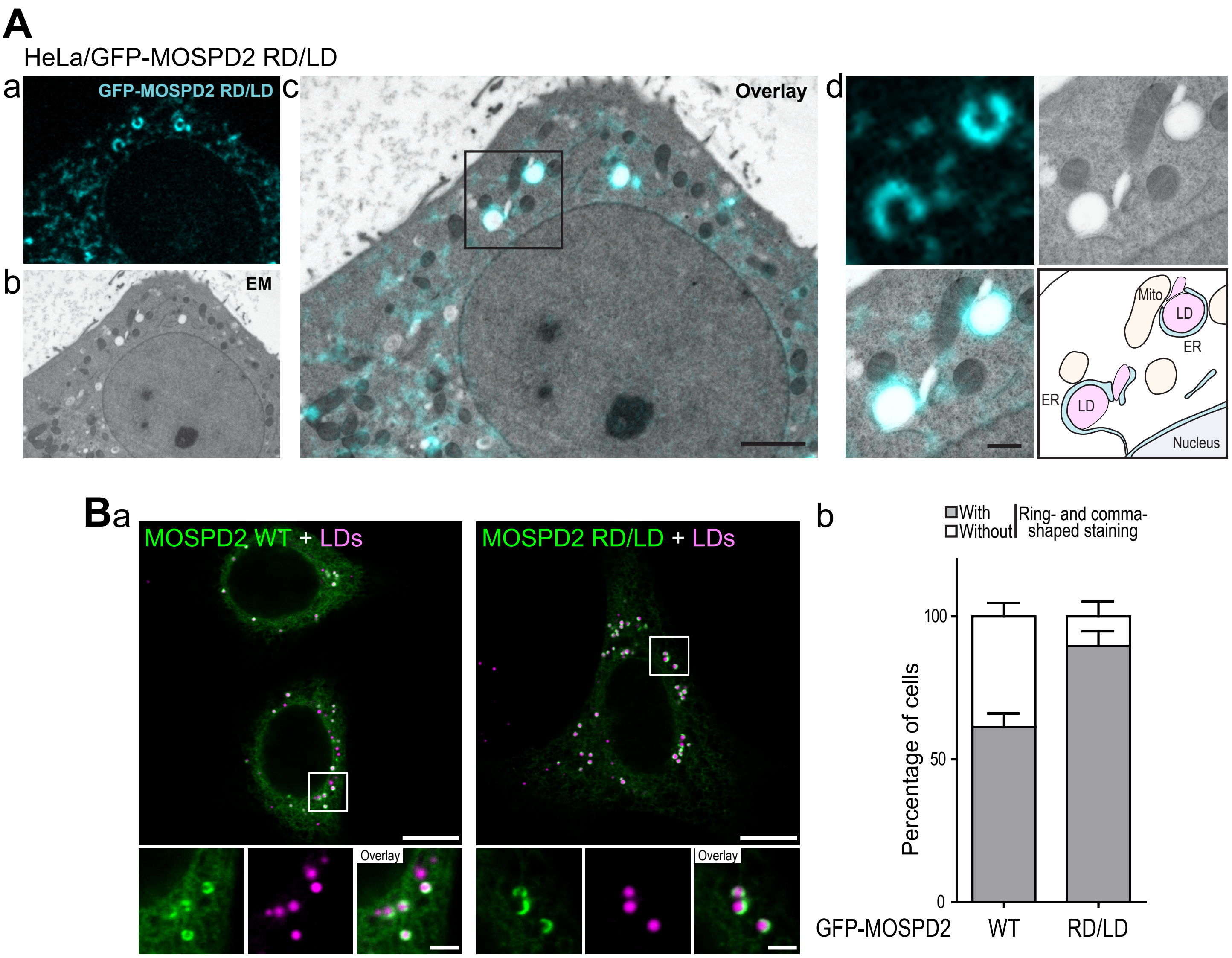


**Figure S7: Subcellular localization of GFP-MOSPD2 RD/LD mutant.**

A.CLEM of a GFP-MOSPD2 RD/LD expressing cell. a: GFP-MOSPD2 RD/LD fluorescence microscopy image; b: EM image; c: correlation of GFP-MOSPD2 RD/LD fluorescence and EM images (scale bar 2 µm); d: higher magnification images of the area outlined in black (scale bar 500 nm); bottom right: interpretation scheme showing contacts between organelles; ER and lipid droplets are in cyan and pink, respectively. Mitochondria, endosomes/lysosomes and nucleus are in yellow, gray and light blue, respectively.

B: a: Representative confocal images of GFP-MOSPD2 WT and RD/LD mutant expressing cells. Cells were not treated with OA. LDs were stained with Nile Red (magenta). Images were acquired on a confocal microscope (Leica SP8; x63 NA 1.4). Scale bar: 10 µm (insets 2 µm). b: percentage of cells with GFP-positive ring- or comma-shaped structures. Mean ± SD; n = 4 independent experiments (WT: 156 cells; RD/LD: 162 cells).

### Figure S8

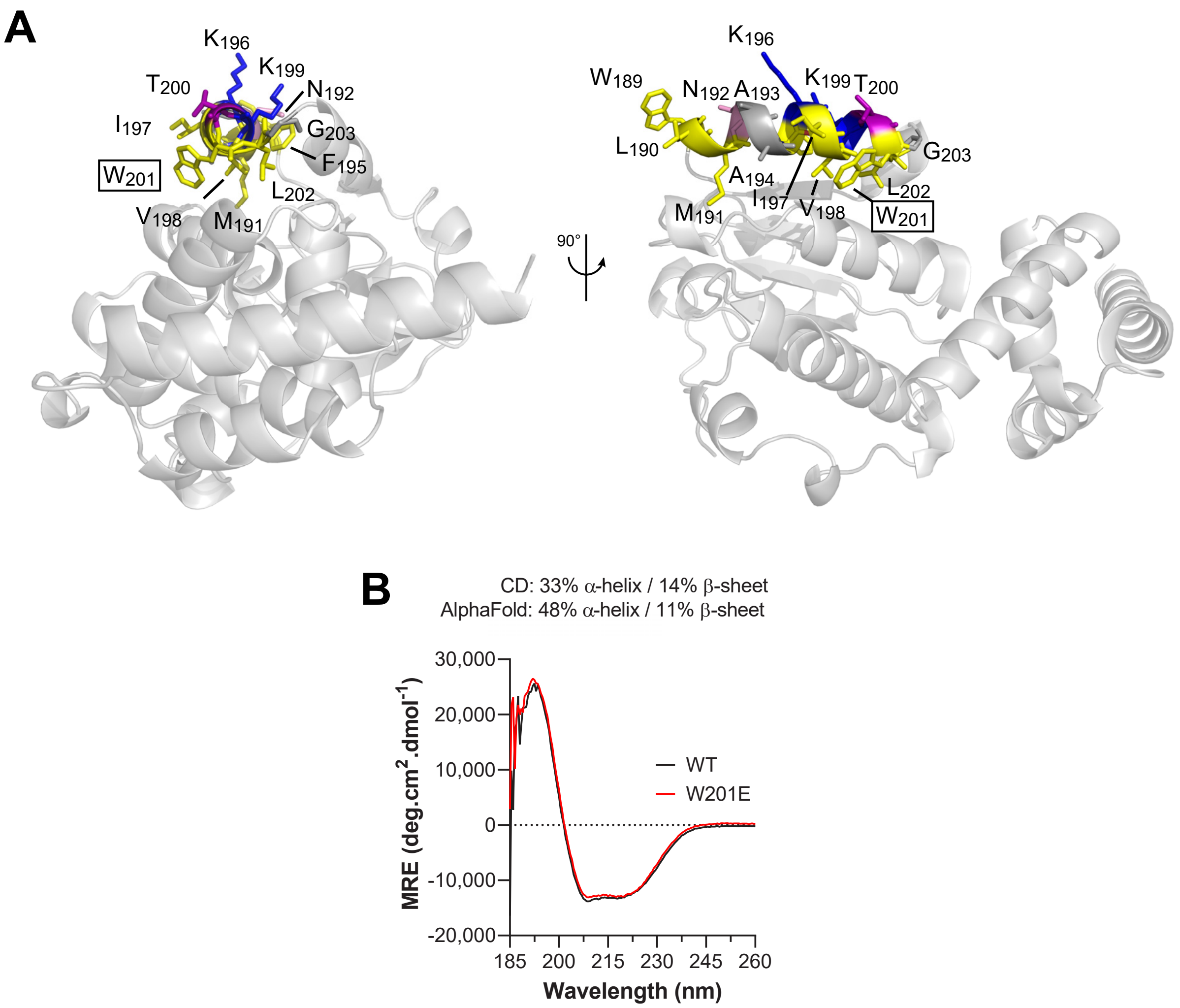


**Figure S8: Structure model of the CRAL-TRIO domain of MOSPD2 and structural assessment.**

A. Ribbon diagram of the structure model of the CRAL-TRIO domain of human MOSPD2 (Uniprot Q8NHP6; residues 1-241) obtained with AlphaFold (Jumper et al, 2021). The domain is in light grey except for the amphipathic helix depicted in stick model with residues colored as in Fig. 5B.

B. Far-UV CD spectrum of the MOSPD2 CRAL-TRIO domain and its W201E variant (6.7 µM) in 20 mM Tris, pH 7.4, 120 mM NaF buffer. The percentage of α-helix, β-sheet and turn, deriving from the analysis of the spectrum (WT) are given as well as the values deriving from the structure model (AlphaFold) using the DSSP algorithm.
